# How Bacterial Growth-Phase Reshapes Synthetic Gene Circuit Function

**DOI:** 10.64898/2026.10.09.752898

**Authors:** Eric Lu, Domitilla Del Vecchio

**Affiliations:** Department of Biological Engineering, MIT, Cambridge, MA, USA; Department of Mechanical Engineering, MIT, Cambridge, MA, USA

## Abstract

The development of biological engineering tools have improved the ability to embed novel functions into living organisms. Concurrently, our understanding of the interface between synthetic genetic circuits and their host has accelerated in the past decade, enabling the development of context-aware predictive models. However, long-term circuit function remains difficult to predict when designs operate in field-relevant conditions. Specifically, while synthetic genetic circuits are typically characterized during exponential growth, these must operate for extended periods of time in nutrient-limited conditions and where bacteria enter stationary growth phase. In this work, we construct a mathematical model that captures the dynamics of a genetic circuit as cells transition into stationary growth phase. To achieve this, we construct synthetic genetic circuits with fluorescent reporter output and measure plasmid DNA copy number, mRNA level, protein level, and live cell count over the course of one week in stationary growth phase. We find that transcription rates plummet shortly after entering stationary phase but settle at a basal value that depends on both the promoter and ribosome binding site. Translation rates also decline in stationary phase, but protein accumulation due to lack of cell division can increase protein levels well beyond exponential phase. Our model recapitulates these findings and is capable of predicting the output of new genetic circuits. This model can serve as a tool to determine how a genetic circuit design may change on longer time scales where cells enter stationary growth phase.

## Introduction

Genetic circuits are typically designed and modeled under lab conditions where bacteria grow exponentially. Accordingly, modeling tools used for genetic circuit design assume exponentially growing cells^1,2,3^. However, many applications of engineered bacteria require long-term deployment in nutrient-limited environments where cells enter stationary growth. For instance, many clinical trials have been conducted to investigate the efficacy of genetically engineered bacteria designed to produce therapeutic molecules in patients’ guts, or modulate the immune system after being intratumorally injected or topically applied, then left on site^4,5,6,7,8^. There has also been growing interest in using engineered bacteria as environmental biosensors to continuously monitor chemical or physical stimuli, as *in situ* bioremediators to remove complex pollutants from the environment, or as “microbial cell factories” to transform renewable carbon sources into chemicals of interest.^9,10,11,12^ All these applications require engineered bacteria to operate in stationary phase where genetic circuits’ behaviors are typically not tested (Fig. 1A).

**Fig. 1.**
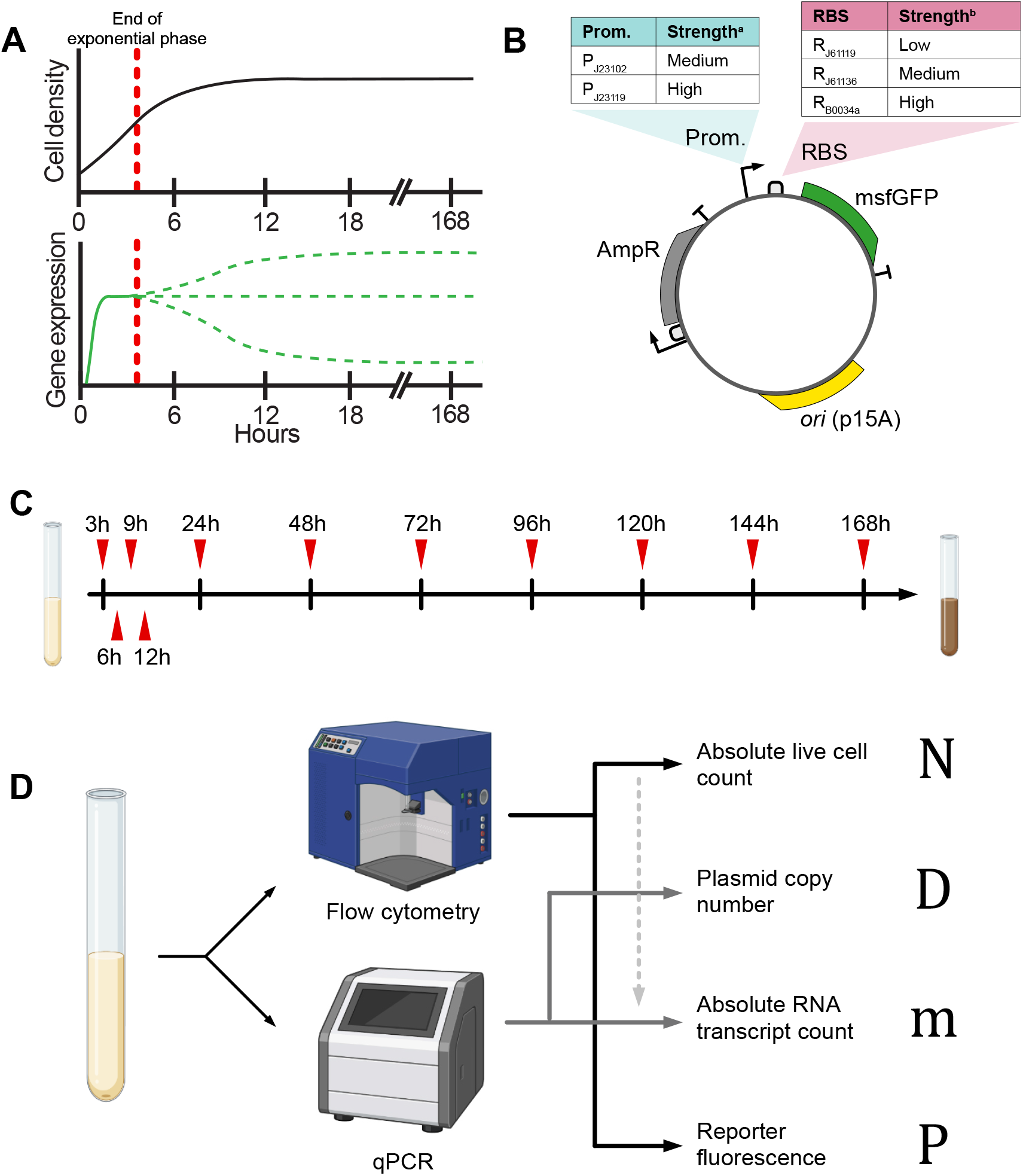
Approach to characterize gene expression dynamics of long-term bacterial cultures. (A) Cell density grows exponentially until nutrient limitations cause it to plateau. Gene expression (green traces) is affected by the transition out of exponential phase. (B) Plasmid encoding the expression system used for characterizing gene expression dynamics. The reporter msfGFP is expressed by different combinations of two promoters and three ribosome binding sites, each with different strengths (Supplementary Table S1). (C) Timeline of measurements, indicated by red ticks (see Methods). (D) Measurement methods. Flow cytometry is used for cell count and protein fluorescence while qPCR is used for plasmid copy number and RNA abundances. RNA abundance is then transformed into absolute values using cell counts (see Methods). ^a^Promoter strength predicted by De Novo DNA’s Promoter Calculator (Version 2022)^21^ ^b^RBS strength predicted by De Novo DNA’s RBS Calculator (Version 2020)^22^

Many studies have quantified changes in the availability or activity of molecular machinery responsible for plasmid replication^13,14,15^, transcription^16,17,18^, and translation^16,19^ between exponential and stationary phase. A study of Klumpp et al. investigated the global influence of growth rate on an exogenous constitutive reporter’s expression by varying the media composition to modulate growth rate and therefore did not examine the natural transition from exponential to stationary phase over time^20^. Moreover, the authors’ model of protein expression were based on plasmid copy numbers and RNA expression that were computed based on other studies rather than on concurrent measurements of those species’ abundance. A characterization of the effects of growth phase on engineered genetic circuits’ behavior remains largely unexplored, but would enable the construction of models of gene expression that can be used in synthetic biology.

In this study, we measure cell concentration, plasmid copy number, transcript abundance, and protein abundance of a plasmid-borne, constitutively-expressed fluorescent reporter over seven days in *E. coli* cells (Fig. 1B). We use these measurements to construct a differential equation model of gene expression that captures the transition from exponential growth phase to stationary phase up to a week. We then test the model’s ability to predict the behavior of gene expression regulation in constructs with different circuits that were not previously used for the fitting of parameters or for model selection.

## Results

### Exponential-to-stationary model of gene expression

To construct an exponential-to-stationary model (ESM) of the dynamics of gene expression of a plasmid-borne gene, we model the temporal change of cell number, plasmid copy number, mRNA level, and protein fluorescence per cell via ordinary differential equations (ODE).

In traditional models of gene expression during exponential growth, plasmid copy number is assumed to be a constant, resulting from the equilibration of plasmid replication with plasmid dilution^1,23^. Similarly, the transcription rate for a constitutively expressed gene is assumed to be a constant, the translation rate is assumed to increase with mRNA concentration, and the decay rates of these species, given by dilution and degradation, are also assumed to be proportional to the species concentrations via a constant dilution rate parameter^1,23^. Transcription and translation rates are also proportional to the concentration of transcriptional and translational resources with the level of these resources often approximated by a constant during exponential growth, which is proportional to the growth rate^1,16,20^. In this paper, we modify this model by accounting explicitly for the dynamics of cell density and for how these dynamics affect transcription, translation, and decay rates. We let *N* denote the cell density and, as a first step to link population-level growth to the production of intracellular species, we let the production rate of each species be proportional to the logistic growth function, 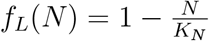 with cell density carrying capacity *K*_*N*_. Furthermore, we explicitly distinguish between degradation of a species and dilution of a species due to cell division, and let the dilution rate be proportional to growth rate. Finally, because translational resources are especially limited in bacteria, we do not assume that protein production is a linear function of mRNA concentration and instead adopt a Michaelis-Menten form for protein production.^24,1^ With these assumptions, and by letting *D* represent the cellular counts of plasmid copy number (PCN), *m* represent the cellular counts of mRNA, *P* represent the fluorescence intensity of the reporter protein, we write the ODEs as:

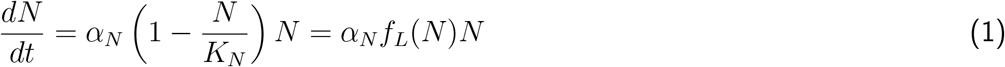

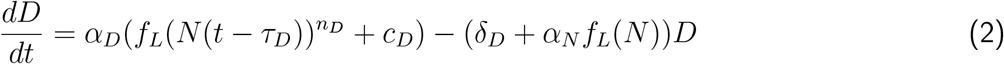

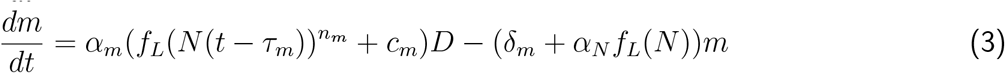

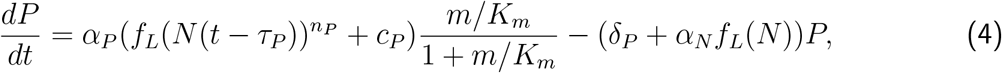

in which *n*_*D*_, *n*_*m*_, and *n*_*P*_ are natural numbers that capture possible cooperative effects of growth rate on production rate^25^, *τ*_*D*_, *τ*_*m*_, and *τ*_*P*_ capture possible delays in the effects of *N* on production rate^26^, and *c*_*D*_, *c*_*m*_, and *c*_*P*_ capture basal production rates that are present even when growth halts.

We estimate the parameters of this model using a step-wise approach. We start by fitting the parameters in Eq. 1 from temporal measurements of cell density *N*. The estimated parameters for Eq. 1 are then fixed, and only the measurements for plasmid copy number *D* are used to estimate the parameters for Eq. 2. Next, estimated parameters for Eq. 2 are fixed (in addition to the fixed parameters for Eq. 1) before measurements of RNA transcript counts *m* are used to estimate parameters for Eq. 3, and similarly for reporter fluorescence *P* in Eq. 4. Model fitting is performed using the ordinary least squares (OLS; Methods). At each step of model fitting, we employ an Akaike information criterion (AIC) to evaluate whether simpler models are more probable, thereby improving model parsimony (Methods).^27^

### Population dynamics

In our model of population dynamics in Eq. 1, the rate of change of *N* (*t*) approaches zero when nutrients are depleted, that is, when *N* = *K*_*N*_. This assumes that cell death is negligible in our experiment, otherwise 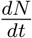 could become zero when the rate of growth equals the rate of death. If this were the case, then our model should include a decay term for *N*, otherwise it would not well-capture how growth rate affects the production of the remaining species in the model. As a first step, we validated the assumption at the basis of Eq 1, that is, that in our experiments the number of dead cells is negligible.

Thus, we asked if the stable cell density after 24 hours was due to a dynamic equilibrium between living and dying cells, or due to a uniformly nongrowing population of live cells. We first quantified the absolute numbers of live and dead cells from 3 to 168 hours post-inoculation using flow cytometry of bacterial cultures prepared with a microsphere standard and propidium iodide (PI, Fig. S1). Live cell counts stabilized at 3.8 *×* 10^9^ cells/mL by 24 hours (Fig. 2A), in line with previous studies performed in similar growth conditions.^28,29^ Moreover, the vast majority of the population was alive (PI-) over a week in batch culture (Fig. 2B).

**Fig. 2.**
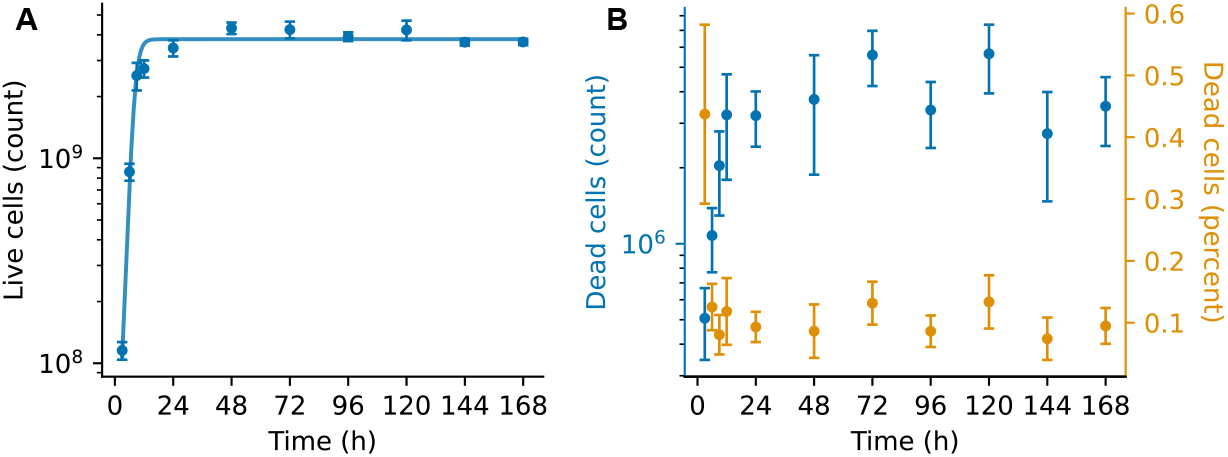
Cell count measurements. (A) Absolute counts of live (PI-) cells over a week in batch culture. Solid line: ordinary least squares (OLS) best fit of Eq. 1 (see Methods; parameters in Table 1). (B) Absolute counts (blue) and percentage of population (orange) of dead (PI+) cells over a week in batch culture.

Then, we reasoned that PI+ cells could still decompose into debris, and therefore that if this decomposition rate is high, then it could play a role in maintaining stationary phase cell concentrations. To rule this out, we estimated the decomposition rate of PI+ cells by treating overnight cultures with 70% ethanol to induce membrane damage, then counted the PI+ population from 0 to 168 hours (Fig. S2). The half-life of PI+ cells was estimated to be 15 hours, confirming that stationary phase is composed of a largely alive but nongrowing population of cells. This therefore validates our assumption and justifies omitting cell death from Eq. (1). Therefore, *N* should be interpreted as both the live cell and total cell population.

**Table 1.** Parameter estimates for live cell counts from Eq. 1.

| Parameter | Value | Unit |
| --- | --- | --- |
| $\alpha_N$ | 0.719 | $\text{h}^{-1}$ |
| $K_N$ | $3.81 \times 10^9$ | Cells $\text{mL}^{-1}$ |

The measurements of cell counts were fitted to the model of population growth (Eq. 1) using ordinary least squares (OLS; Methods). Values for the specific growth rate *α*_*n*_ and carrying capacity *K*_*N*_ are reported in Table 1.

### Plasmid copy number dynamics

To estimate the parameters for the PCN dynamics in Eq. 2, we measured the PCN over time for the p15A and pSC101 origins of replication. We observed that the PCN temporal profile for the p15A origin is biphasic, increasing from 3 to 8 copies in the first 18 hours, then declining back to 3 copies over the rest of the week (Fig. 3B). These values are consistent with those found in previous studies measuring p15A copy number in nutrient rich media.^30,31^

**Fig. 3.**
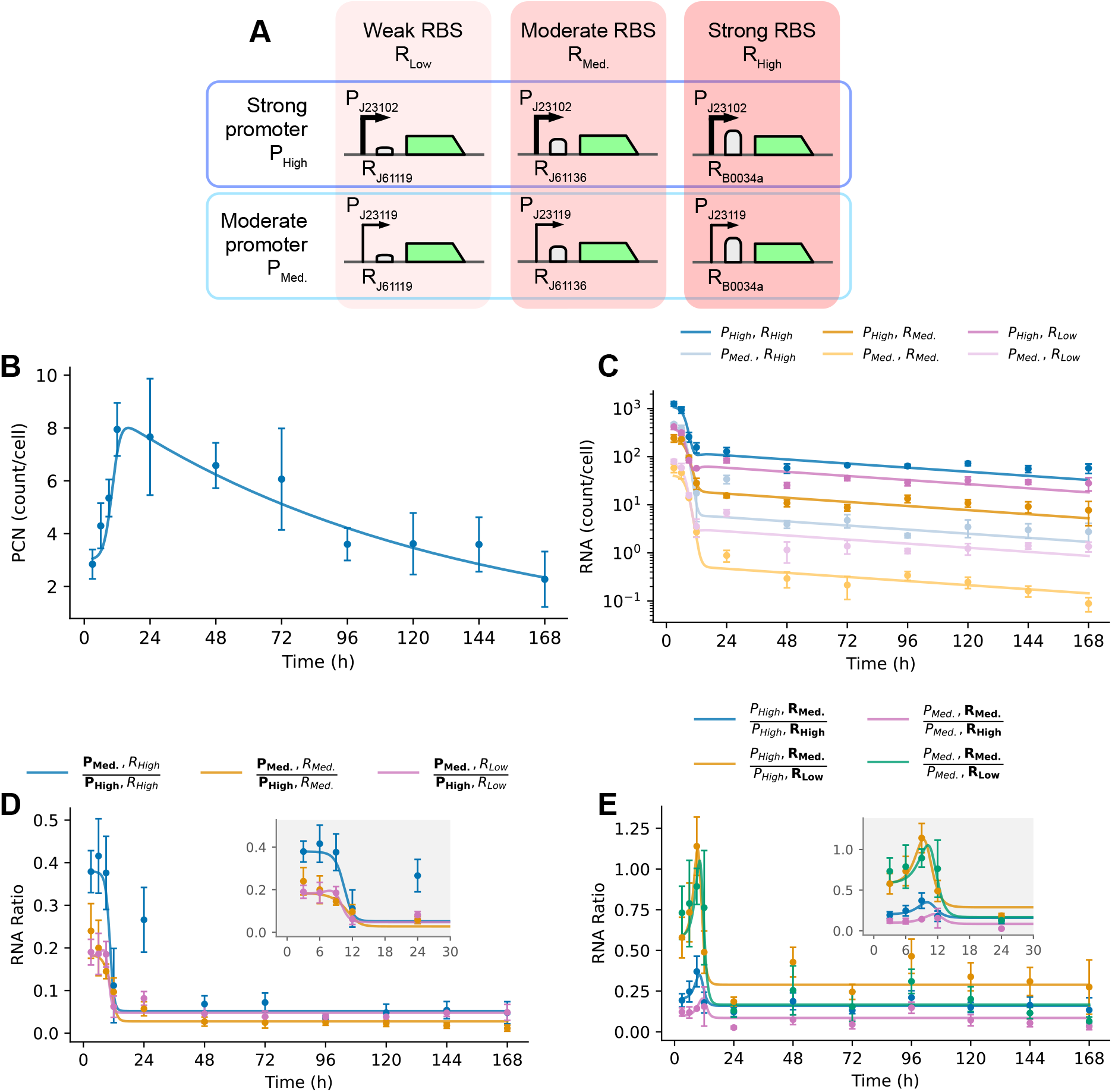
Measurements of intracellular molecular species. (A) Constructs encoding the expression systems used to characterize different promoters and RBSs. (B) Plasmid copy number over time (p15A ori). Solid line: OLS best fit of Eq. 1 and Eq. 5 based on the data shown (see Methods; parameters in Table 2). (C) Absolute RNA transcript counts per cell over time for all six constructs with pairings between two promoters (medium and high strengths) and three ribosome binding sites (RBSs) (low, medium, and high strengths). The notation for each component is listed in Fig. 1B. Solid lines: simulation of the OLS best fit of Eq. 1 and Eq. 5 to 7 based on the data shown (see Methods; parameters in Table 3). (D) Ratio of RNA counts between medium-strength promoter *P*_*Med*._ (numerator) and the high-strength promoter *P*_*High*_ (denominator) for each RBS: *R*_*High*_ (blue), *R*_*Med*._ (orange), *R*_*Low*_ (pink). Comparison groups are bolded in the legend for clarity. Inset: zoom-in of the 0 to 30 h window. Solid lines: transcript ratios based on simulated OLS best fits seen in panel B. (E) Ratio of RNA counts between constructs harboring the medium-strength RBS *R*_*Med*._ (numerator) and each other construct as the denominator: *P*_*High*_ with high-strength RBS *R*_*High*_ (blue), *P*_*High*_ with low-strength RBS *R*_*Low*_ (orange), *P*_*Med*._ with high-strength RBS *R*_*High*_ (pink), *P*_*Med*._ with low-strength RBS *R*_*Low*_ (green). Comparison groups are bolded in the legend for clarity. Solid lines: transcript ratios based on simulated OLS best fits seen in panel B.

We identified a similar increase in plasmid copy number for the pSC101 origin of replication (Fig. S3). However, while p15A copy number declines throughout late stationary phase, pSC101 copy number remains more steady, potentially consistent with the fact that the p15A origin of replication relies on noncoding RNA, whose abundances are especially dynamic across growth phases, while the pSC101 origin relies on proteins to replicate.^32,33,34^

**Table 2.**
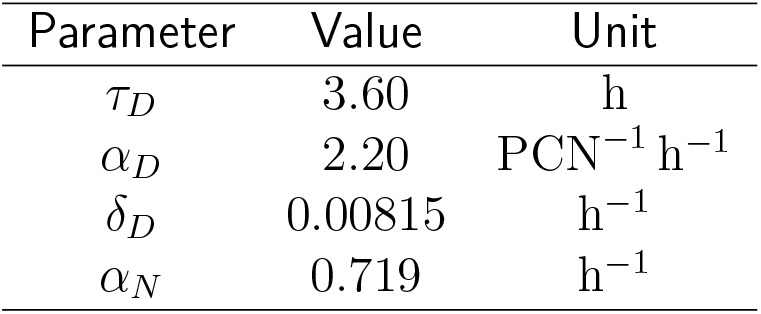
Parameter estimates associated with the p15A replication origin from the PCN model in Eq. 5.

**Table 3.** OLS parameter estimates for each Promoter and RBS combination associated with the RNA model in Eq. 7 (*α*_*m*_ in *RNA/*(*PCN* · *h*); transcription rates in *RNA/h*). Exponential phase (EP TR) and stationary phase transcription rates (SP TR) were computed by isolating the production rate term in Eq. 7, normalizing by *D*, and taking the extreme values of *f*_*L*_(*N*) for each of the listed parameter sets. *δ*_*m*_ = 8.32*h*^−1^ (5 minute half-life) used for each combination.^38^

| Promoter | RBS | $n_m$ | $c_m$ | $\alpha_m$ | EP TR | SP TR |
| --- | --- | --- | --- | --- | --- | --- |
| $P_{J23102}$ | $R_{J61136}$ | 1.45 | 0.00420 | 122 | 123 | 0.513 |
| $P_{J23119}$ | $R_{J61136}$ | 1.31 | 0.0286 | 651 | 669 | 18.6 |
| $P_{J23102}$ | $R_{B0034a}$ | 1.82 | 0.00482 | 1250 | 1260 | 6.02 |
| $P_{J23119}$ | $R_{B0034a}$ | 1.87 | 0.0366 | 3190 | 3310 | 117 |
| $P_{J23102}$ | $R_{J61119}$ | 1.94 | 0.0149 | 206 | 210 | 3.07 |
| $P_{J23119}$ | $R_{J61119}$ | 2.28 | 0.0583 | 1100 | 1170 | 64.4 |

The cell count-associated parameters in Eq. 1 were fixed before estimating parameters for PCN in Eq. 2 using the measured PCN values at each timepoint and the *N* (*t*) trajectories generated by Eq. 1. However, because we were unable to find estimates for plasmid decay rates in stationary phase bacteria from the literature, we estimated *δ*_*D*_ = 0.00815*h*^−1^ based on the observed decrease in PCN from the p15A origin of replication after 24 hours in culture. To do so, we assumed that plasmid replication rate was zero because the p15A origin of replication relies on RNA to maintain its copy number and RNA transcription rates are known to be reduced in stationary phase (Fig. 3C).^16^

We then performed AIC-based model selection where models of the form of Eq. 2 were considered, but with different parameters omitted (Table *S*6). The model with the lowest AIC had both the cooperativity constant *n*_*D*_ and basal production constant *c*_*D*_ omitted. Omission of *c*_*D*_ is consistent with our earlier assumption made to estimate the decay rate *δ*_*D*_ that the replication rate for the p15A origin of replication is zero in stationary phase. Thus, the resulting model for the p15A origin of replication is:

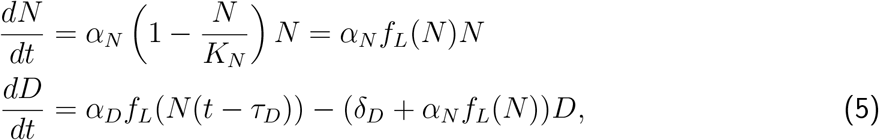

with parameter values given in Table 2. See Table S4 for parameter estimates of the pSC101 origin of replication. Similar to how the cell population parameters were fixed to estimate parameters of and select a parsimonious model for PCN dynamics, this reduced PCN model is then used in subsequent model fitting routines for modeling RNA and protein abundance.

### Per-cell transcript number dynamics

With a model describing both the cell density and PCN dynamics over time we can now estimate the parameters of the ODE of transcript number using temporal mRNA count measurements. To measure the reporter gene RNA copy number per cell, we first generated a standard curve using known amounts of reporter mRNA (Fig. S4). Then, we quantified transcript abundance in our experiments using RT-qPCR (see Methods). Finally, we normalized the number of transcripts by the number of cells used for RNA extraction and computed the average number of transcripts per cell (Methods).

We quantified absolute transcript counts over time for six reporter constructs harboring combinations of two promoters and three RBSs with different strengths as predicted by transcription and translation initiation rates calculators, respectively (Fig. 3A).^21,22^ Because the RBS sequence can influence transcript counts by modifying, for example, the rate of transcription initiation or the transcript’s stability, a unique set of parameters for Eq. 3 was estimated for each promoter and RBS combination.^35^

We observed that transcript counts for each combination are constant over time between 3 and 6 hours post-inoculation, but decline sharply to a range of 3.6% to 13.7% of exponential phase abundances by 12 hours, ranging based on the promoter and RBS combination (Fig. 3C). By 1 week in stationary phase, counts fall further to a range of 0.15% to 6.7%, ranging based on the promoter and RBS combination, of exponential phase counts with the later decline concomitant with falling PCN (Fig. 3B).

Many genetic circuits only generate a desired behavior when their component parts operate within specific ranges of relative expression levels.^36^ Thus, we investigated if the relative transcript abundance between constructs, which we call RNA ratio, are maintained throughout growth phases by first comparing transcript abundance between two promoters paired with the same RBS (Fig. 3D). At the 3 hour timepoint, the moderate-strength P_J23102_ promoter expresses between 18% and 38% (varying based on RBS) of the strong P_J23119_ promoter’s transcripts. This ratio remains steady for 9 hours before declining sharply.

Similarly, a comparison of RNA ratios across the three RBSs paired with the same promoter reveals that RNA ratios fluctuate over the first 24 hours before stabilizing (Fig. 3E). Thus, both RBS and promoter choice can substantially alter how RNA ratios fluctuate across and throughout growth phases. This suggests that circuits tested in one growth phase may fail when operating in another phase or across multiple phases. Ratios plotted with transcript abundances of the RBSs *R*_*J*61136_ and *R*_*B*0034*a*_ as the numerator are available in Fig. S5.

Parameters for the transcript dynamics in Eq. (3) were then estimated for each of the six constructs, followed by AIC-based model selection where models with different parameters omitted from Eq. 3 were fit on the transcript count trajectory over time and the AIC for each model was computed (Table S6). We assumed a constant nominal degradation rate *δ*_*m*_ = 8.32*h*^−1^ based on available literature^37^ and included an additional constant 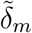 to capture sequence-specific variability in mRNA degradation rates that is estimated alongside the other parameters during AIC-based model selection to yield the following differential equation for mRNA dynamics:

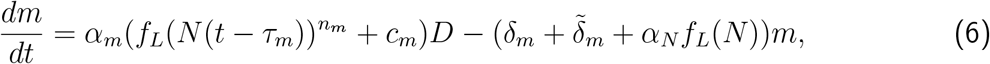

where *δ*_*m*_ = 8.32*h*^−1^ is fixed and 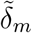 is estimated using OLS. We also note that previous studies have found that mRNA half-life is higher in stationary phase than in exponential phase, implying that our parameters overestimate transcript production rates in stationary phase since 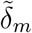 is a constant.^38,39^

Models with 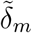 omitted had lower AICs than those including 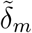 for each set of parameters omitted from Eq. 6 (Table S6). Thus, flexibility for sequence-specific mRNA degradation rates can be omitted in favor of model parsimony. While models with both *τ*_*m*_ and *n*_*m*_ omitted from Eq. 3 had the lowest AIC for 3 of the 6 constructs, models with the delay term *τ*_*m*_ omitted had lower AICs than the full model across all 6 constructs (Table S6). Thus, we opted to only omit *τ*_*m*_ in the model to represent transcript dynamics:

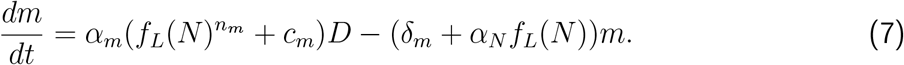

The fitted model yielded exponential phase transcription rates, derived from Eq. 7 as *rate* = *α*_*m*_(1 + *c*_*m*_) by setting *N* (*t* − *τ*_*m*_) = 0, ranging from 122.6 to 3306.6 RNA*/*(PCN · h), and stationary phase transcription rates, derived from Eq. 7 as *rate* = *α*_*m*_*c*_*m*_ by setting *N* (*t* − *τ*_*m*_) = *K*_*N*_, between 0.5 and 116.6 RNA*/*(PCN · h) (Table 3).

### Protein expression dynamics

With the models identified for cell density, PCN, and RNA count dynamics, we now model the dynamics of protein expression using flow cytometry timecourse data. Flow cytometry measurements of msfGFP fluorescence reveal steady fluorescence over time for constructs harboring RBSs *R*_*B*0034*a*_ and *R*_*J*61119_, and increasing protein abundance for constructs harboring RBS *R*_*J*61136_ in the first 24 hours (Fig. 4A). Protein levels for most constructs peak around 48 hours. Thus, while mRNA transcripts are most abundant in exponential phase, protein abundance in stationary phase (at 48 h) is 1.04 to 4.08 times greater than protein abundance in exponential phase (3 h; Fig. 4B). Thus, despite substantial transcription rate decrease, high protein stability and low protein dilution rate can enable protein output to continue increasing in stationary phase.

**Fig. 4.**
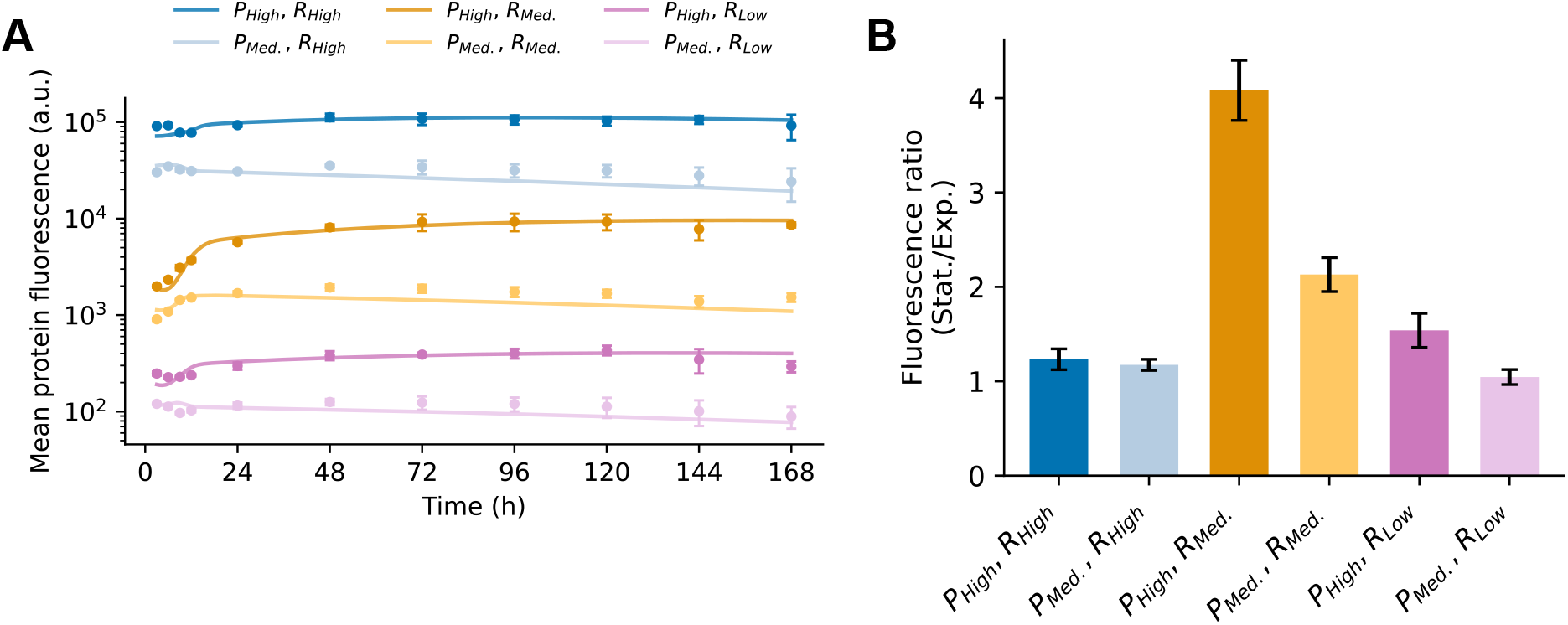
(A) Mean protein fluorescence over time for all six constructs. Solid lines: simulation of the OLS best fit of Eq. 8 based on the data shown (see Methods; parameters in Table 4). (B) Fold change of protein fluorescence at stationary phase (48h, numerator) compared to exponential phase (3h, denominator). Data points in each panel represent the mean ± one standard deviation calculated from three biological replicates.

Before performing OLS fits of the parameters in the protein dynamics ODE (Eq. 4), the msfGFP degradation rate constant *δ*_*P*_ was estimated as 4.2 *×* 10^−3^*h*^−1^ in a separate experiment by inducing msfGFP expression for 24 hours, then washing the inducer away and monitoring cells for fluorescence decay in inducer-free media over the following week (Fig. S6). Since dilution due to cell division is negligible after 24 hours, our decay estimate is an estimate of the degradation rate (Fig. 1A).

**Table 4.** OLS parameter estimates for each RBS associated with the protein model in Eq. 8. *δ*_*P*_ = 4.2 *×* 10^−3^*h*^−1^ and *α*_*N*_ = 0.719*h*^−1^ used for each RBS.

| RBS | $\tau_P$ (h) | $\alpha_P$ (a.u.RNA<sup>-1</sup> h<sup>-1</sup>) | $K_m$ (RNA) | $c_P$ |
| --- | --- | --- | --- | --- |
| $R_{J61136}$ | 7.78 | 1130 | 30.6 | 0.214 |
| $R_{J61119}$ | 2.99 | 141 | 52.5 | 0.0382 |
| $R_{B0034a}$ | 5.80 | $1.11 \times 10^5$ | $1.56 \times 10^3$ | 0.114 |

We then performed OLS fits of the parameters in the protein dynamics ODE using the measured fluorescence values over time with the parameters in the cell count, PCN, and mRNA count dynamics fixed to the values identified before. For each RBS, we performed a different OLS fit for the parameters *α*_*P*_, *τ*_*P*_, and *n*_*P*_, and kept these same parameter values for the different promoters (using the promoter-specific parameters as in Table 3). This assumes that the protein translation rate of a transcript is not directly influenced by the promoter, which is motivated by the fact that the promoter is not transcribed into the mRNA. Each of our constructs is designed without 5’ UTRs, which prior studies have noted can affect translation rate.^40^

Each fit was again followed by AIC-based model selection where models with different parameters omitted from Eq. 4 were fit on the fluorescence data and the AIC for each model was computed (Table S6). While models omitting either *n*_*P*_ or *τ*_*P*_ resulted in AIC values lower than the full model’s AIC for RBSs *R*_*J*61119_ and *R*_*J*61136_, only the model omitting *n*_*P*_ had a lower AIC than the full model’s. Thus, for simplicity, we select the protein level differential equation that omits *n*_*P*_, resulting in the following differential equation:

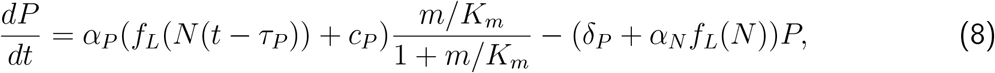

with parameter values as in Table 4.

The rate at which the fluorescence intensity increases varies across several orders of magnitude across the different RBSs, with *R*_*B*0034*a*_ driving the fastest increase and *R*_*J*61119_ the slowest increase of protein abundance. These observations align with the strength (transcription initiation rate) of each RBS as predicted using a transcription initiation calculator (Figure 1A).^22^

The constant *K*_*m*_ tunes the saturation of the translation rate when the mRNA level increases, depends on the specific RBS, and varies modestly between 30.6 and 52.5 counts for the RBSs *R*_*J*61119_ and *R*_*J*61136_, respectively, but is 1562.2 counts for *R*_*B*0034*a*_.

The delay constant *τ*_*P*_ also varies across the different RBSs, from 3.0 to 7.8 *h*. Nonetheless, the protein expression rate decreases by 78.6% to 96.2% of exponential phase rates in deep stationary phase, as indicated by the estimates for constant *c*_*P*_.

### Model Validation

We then assessed the predictive ability of the ESM, focusing on how its circuit-independent parameters and architecture estimated previously is predictive of outputs from new circuit topologies. These circuit-independent aspects are: the estimated cell density and PCN dynamics, and the final structure of the differential equations describing the mRNA and protein dynamics, specifically, the omission of *τ*_*m*_ in Eq. 7 and the omission of *n*_*P*_ in Eq. 8. We used the ESM to predict protein output over a week for different constitutive promoters not used before and for both inducible promoters. Specifically to test different constitutive promoters, we kept all the parameter values unchanged from the prior estimates except for those pertaining to the promoter itself (*α*_*m*_, *c*_*m*_, and *n*_*m*_; while *τ*_*m*_ was omitted as previously). First, we tested the model’s ability to predict the output for different promoters by generating 18 more constructs harboring different promoters from the Anderson library^41^, each paired with the *R*_*J*61136_ RBS (Fig. 5A).

**Fig. 5.**
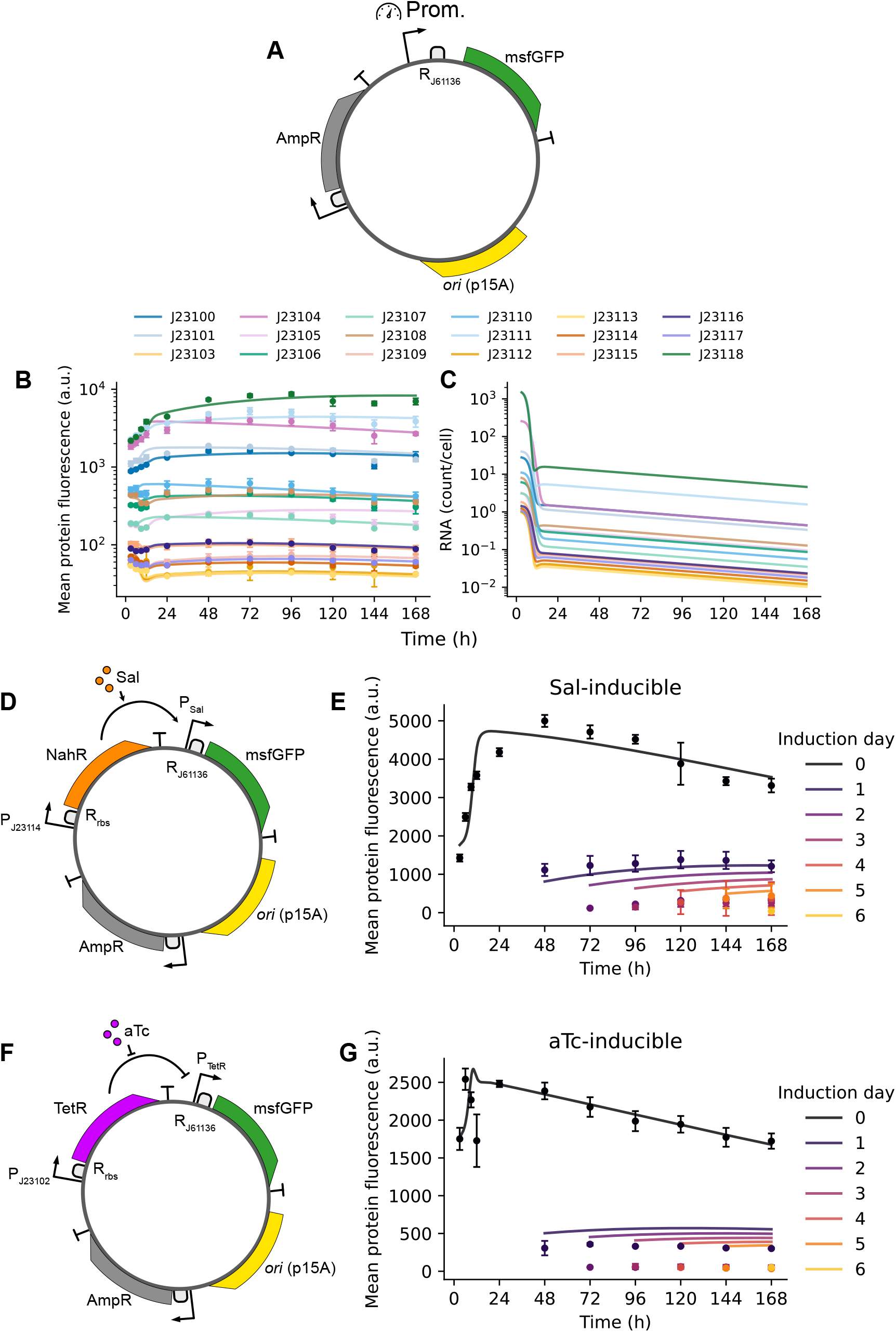
Model validation. (A) Plasmid map of the expression system used for model validation with 18 promoters from the Anderson library. All systems use the R_J61136_ RBS and promoters following the P_J231–_ naming convention. (B) Mean protein fluorescence over time (24 to 168 h) for 18 promoters from the Anderson library, all paired with the moderate-strength RBS *R*_*J*61136_. Solid lines: simulation of the OLS best fit of Eq. 1 and Eq. 5 to 8 with parameter values given in Table S5 (See methods). Specifically, only *α*_*m*_, *c*_*m*_, and *n*_*m*_ pertaining to promoter choice were fitted while all other parameters were unchanged. (D) Plasmid map of the Sal-inducible circuit. (E) Protein fluorescence over time of cultures harboring the Sal-inducible circuit induced with 1 mM Sal from 0 to 6 days in 24 h intervals. Each color corresponds to the day that a separate set of cultures was induced with Sal. Solid lines: simulations of Eq. 1 and Eq. 5 to 14 with *I* set to 1000 *a*.*u*. from 0 to 6 days in 24 h intervals. See Table S3 for full parameter set used in simulations. F) Plasmid map of the aTc-inducible circuit. (G) Protein fluorescence over time of cultures harboring the aTc-inducible circuit induced with 1 *µ*M aTc from 0 to 6 days in 24 h intervals. Solid lines: simulations of Equations 1, 5 to 8, and 17 to 20 with *I* set to 1000 *a*.*u*. from 0 to 6 days in 24 h intervals. Data points in panels A, D, and F represent the mean ± one standard deviation calculated from three biological replicates.

To this end, we measured protein fluorescence for each construct from 24 *h* to 168 *h* in culture (Fig. 5B). Then, we estimated only the parameters *α*_*m*_, *c*_*m*_, and *n*_*m*_ from the temporal fluorescence data by fixing the previously-estimated parameters for cell density and PCN as in Tables 1–2 and the protein fluorescence parameters for *R*_*J*61136_ as in Table 4. Thus, the transcription rate constant *α*_*m*_, basal constant *c*_*m*_, and exponent *n*_*m*_ are the only remaining free variables for Eq. 7. The resulting predictions from the ESM, in which the RNA differential equation has parameters given by the new estimates, well-describe the temporal trajectories of the proteins in the new promoter configurations (Fig. 5A). We also show that even if mRNA counts are not measured, the ESM also produces mRNA concentration temporal profiles (FIg 5C), which can be used to aid circuit design. (Fig. 5C).

We finally sought to determine if the ESM could recapitulate the behavior of a circuit involving more than one exogenously produced gene, and in a configuration involving transcriptional regulation. To this end, we built two small-molecule inducible msfGFP expression systems, both using the p15A origin of replication plasmid. The first circuit constitutively expresses NahR, a regulator protein that binds the P_Sal_ promoter in the presence of sodium salicylate (Sal) to activate msfGFP expression (Fig. 5D). The second system constitutively expresses TetR which binds the P_TetR_ promoter, repressing msfGFP expression, unless bound to anhydrous tetracycline (aTc) (Fig. 5F).

We then induced batch cultures for each construct every 24 hours from 0 h to 144 h. The induction at 0 hours represents exponential phase induction. As expected, the greatest difference between two induction timepoints was observed between 0 h and 24 h likely due to the large decrease in transcription rate by 12 hours across all previously characterized constructs (Fig. 5E, G). While induction at 24 h still yielded protein expression, cells were mostly unresponsive to induction by 48 hours.

We adapted the ESM to include the mRNA and protein for both the regulator protein (NahR or TetR) and msfGFP, the interaction of the regulator with its cognate small molecule, and the binding of the regulator with/without small molecule to the promoter of msfGFP, following the models of induced transcriptional regulation described in [1]. With differential equations describing cell population and plasmid copy number kept the same as before (Eq. 1 and Eq. 5), the system describing the Sal-activated circuit is:

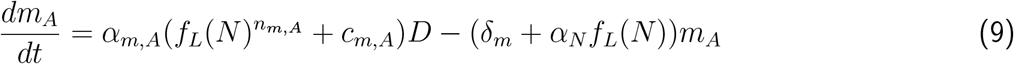

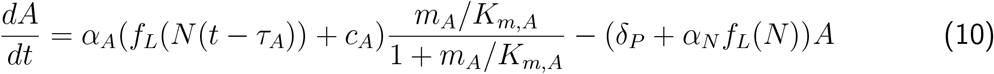

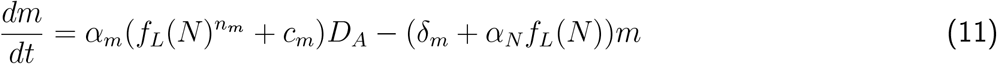

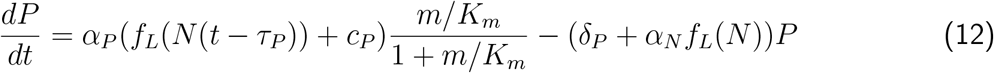

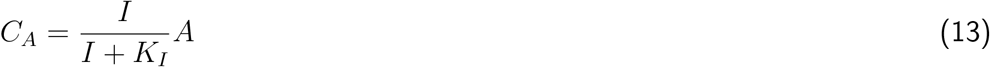

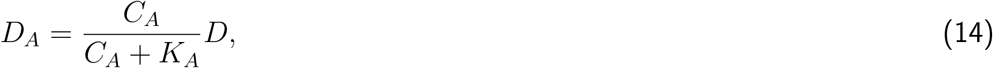

constant for *C*_*A*_ binding to the reporter’s promoter, and *D*_*A*_ represents the complex of *C*_*A*_ bound to the promoter driving the production of *m*. These expressions take the Michaelis-Menten form obtained by applying the appropriate conservation laws and assuming that the complexes *C*_*A*_ and *D*_*A*_ have reached quasi-steady state (See Supplementary Note).^1^ Similarly, the system describing the aTc-activated (de-repressed) circuit is:

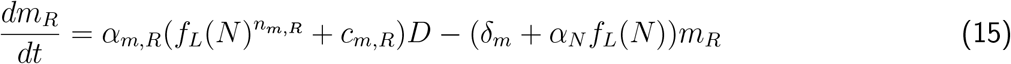

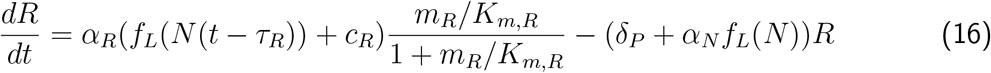

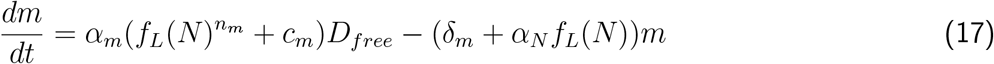

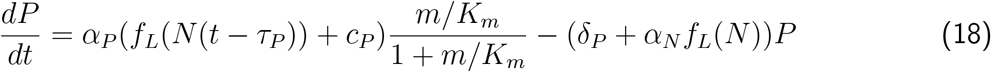

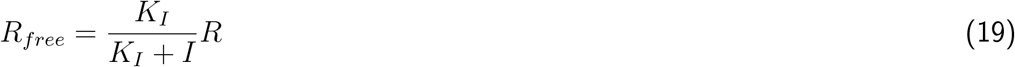

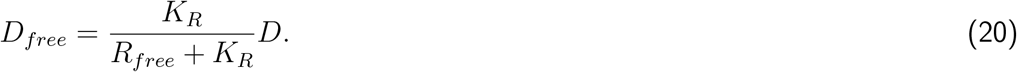

in which *R*_*free*_ represents the amount of TetR not bound to aTc (I), *K*_*R*_ is the dissociation constant for *R* binding to the reporter’s promoter, and *D*_*free*_ represents the promoter that is not bound by TetR, driving the production of *m. R*_*free*_ and *D*_*free*_ are derived using a similar process as *C*_*A*_ and *D*_*A*_ in Eq. 13–14 (See Supplementary Note).

In both systems, we further assume that *I* ≫ *K*_*I*_ to reflect that we use saturating inducer amounts. Therefore, in the above models we have that *C*_*A*_ ≈ *A* and *R*_*free*_ ≈ 0 and thus precise values for *I* and *K*_*I*_ are not necessary.^42^ The degradation rates of all protein species are assumed to be equal and given by the value *δ*_*P*_ = 4.2 *×* 10^−3^*h*^−1^, as estimated previously. Similarly, the degradation rates of all mRNA species are assumed to be equal and given by the value *δ*_*m*_ = 8.32*h*^−1^, as set before.

Parameters for population growth and PCN dynamics were fixed as in Tables 1 to 2. Parameters for NahR (Eq. 9, 10) and TetR (Eq. 15, 16) transcription and translation dynamics were fixed to those previously fitted for P_J23102_:R_B0034a_ based on their predicted transcription and translation initiation rates (see Tables 3 and 4 and Table S2). Inducible msfGFP’s RBS was R_J61136_, thus the RBS-associated parameters for both Eq. 12 and Eq. 18 were fixed as in Table 4. The remaining free parameters describe the transcription of the inducible promoters (*α*_*m*_, *c*_*m*_, and *n*_*m*_) for both equations 11 and 17, and the dissociation constants for *C*_*A*_ and *R*_*free*_ binding to DNA (*K*_*A*_ and *K*_*R*_). The free parameters were estimated using only the protein fluorescence measurements from the 0-hour induction condition. The full list of estimated parameter values used for simulations are provided in Table S3.

Simulations of the models in equations 9–20 under different induction conditions, wherein the time of induction is changed within the first 6 days, return a protein expression profile well-representing experimental data despite parameters being estimated by only using temporal fluorescence data for the samples induced at 0 hours. In particular, the model predicts that later inductions result in lower protein expression, consistent with the data, although slightly overestimating the protein level for inductions occurring after 24 hours. (Fig. 5E, G).

Finally, we performed fluorescence measurements of the P_J23102_:R_J61136_ and P_J23119_:R_J61136_ promoter-RBS pairs after swapping the p15A replication origin for pSC101 to determine whether the fluorescence output could be predicted using the ESM after only changing PCN-associated parameters in Eq. 2. However, the predictions overestimated actual measurements by roughly a factor of two across all timepoints (Fig. S7). We hypothesize that this is because the pSC101 origin of replication also encodes for the repA gene which was positioned immediately downstream of the msfGFP gene, while the p15A origin of replication does not encode another protein. This configuration between two nearby genes is known to reduce both genes’ expression through supercoiling effects.^43^

## Discussion

While new genetic circuits are customarily tested and modeled in exponentially growing cells, how such circuits may function when cells reach stationary growth phase has remained elusive; yet, many applications of engineered bacteria require long-term deployment in growth-limiting conditions. Our study uncovers how growth phase influences exogenous genes’ expression by quantifying changes in the dynamics of processes constituting the central dogma of molecular biology.

We began by investigating whether cells in the stationary phase are constantly growing and dying, or simply not growing. Some studies refer to a “death phase” when 90 – 99% of a bacterial population “die” based on their inability to produce colony-forming units after approximately 3 days of continuous culture in permissive growth conditions.^44^ We found that over 99% of bacteria are still intact and express genes at low levels up to a week in stationary phase (Fig. 2B), and therefore for our application these cells are to be considered alive. Our use of PI staining enabled us to identify that most bacteria in stationary phase are intact and “living”, regardless of their colony-forming potential. Thus, we modeled bacteria as a homogeneous population throughout stationary phase (Eq. 1).

Our results also confirm that plasmid copy number is an important source of gene expression variability.^45^ From exponential to early stationary phase, copy numbers of both p15A and pSC101 replication origins increase sizably (Fig. 3B), though different origins of replication may have different longer-term behaviors due to their distinct mechanisms for replication and copy number maintenance (Fig. S3).

We found that transcription rates strongly attenuate after 6 hours in culture, and both the choice of promoter and RBS can have large effects on the change in transcript abundance in the transition from exponential to stationary phase. In the case of the constructs *P*_*High*_, *R*_*Low*_ and *P*_*Med*._, *R*_*High*_, both had comparable transcript abundances in exponential phase, but deviated substantially at 12 hours in culture (Fig. 3C). Differences in transcript abundances were greatest in stationary phase when comparing between constructs (Fig. 3D, E). Thus, genetic circuits’ behaviors, especially circuits employing functional RNAs beyond protein synthesis, are likely to change substantially across growth phases.

In contrast to transcript abundances, protein abundance for msfGFP remains relatively steady and peaks during stationary phase (Fig. 4A, B). Our model attributes this to the long half-life of msfGFP and reduced rate of protein dilution as cell growth slows down. Low levels of transcription and translation between 12 hours and 48 hours then cause a slow but steady accumulation of protein in stationary phase – up to over four-fold higher than in exponential phase (3h) with one of the tested constructs (Fig. 4A, B). Taken together with the changes in transcript levels, these results suggest that transitioning between growth phases can disrupt behaviors of genetic circuits tuned in exponential phase.

Experimentally, we focused on only the influence of growth phase on gene expression by investigating the behavior of simple constructs within a well-known bacterial strain and fixed growth conditions. However, additional bacterial strains and growth conditions are necessary to understand how other environmental variables influence gene expression to predict circuit function for field deployment.

Additionally, during our validation studies, the ESM overestimated the amount of induced gene expression after 2 days of induction (Fig. 5E, G). This is consistent with known physiological changes where *E. coli* reduce their cell membrane permeability in stationary phase growth.^46,47^

As a phenomenological model, the ESM is simple enough to be used for the design of genetic circuits. As such, the cooperativity coefficients and delay differential equations used in this study were used to simplify models rather than characterize the dynamics of any of the underlying molecular mechanisms. Additionally, the decay rate constants, *δ*, were not optimized within this study. Thus, true variations in the decay rates of species would instead appear as changes to the parameters that were estimated. We also found large variations in the delay constant *τ*_*P*_ across RBSs (Table 4), potentially due to the effects of translational resource partitioning where a decrease in translation rate is delayed with respect to the slowing of population growth by reallocating ribosomes from translating growth-related transcripts to translating transcripts from exogenous genes.^48^

Nonetheless, this simplistic representation of system dynamics ESM backed by experimental results offers a simple conceptual framework for making predictions on synthetic circuit behavior across growth phases, anticipating potential failure modes in circuits designed for long-term deployment, and reducing the time and labor required to test circuits in the lab.

## Acknowledgements

We thank Dr. Ray Henson for useful discussions. This research was developed with funding from the Defense Advanced Research Projects Agency (DARPA) under Contract HR001124C0441, and with partial funding by the Vannevar-Bush Faculty Fellowship (Office of Naval Research award no. N00014-25-1-2053). The views, opinions and/or findings expressed are those of the author(s) and should not be interpreted as representing the official views or policies of DARPA, the Department of War, or the U.S. Government. Approved for Public Release, Distribution Unlimited.

## Methods

### Bacterial culture conditions

*E. coli* MG1655 were grown in 3 mL of M9 minimal media in 14 mL polystyrene culture tubes at 30°C, continuously shaking at 250 rpm. The experiment’s growth medium is an M9 minimal medium composed of M9 salts (1X), 0.4% glucose (SIGMA-ALDRICH, 49159), 0.2% casamino acid (VWR, TS61204-5000), and 1 mM thiamine hydrochloride (SIGMAALDRICH, T4625-25G) supplemented by 100 *µ*g/mL ampicillin. Each culture was briefly vortexed prior to measurement.

### DNA Construction

The genetic circuit construction was based on Gibson assembly methods using pKMDDV_Plasmid_1 (Addgene plasmid ID: 249037) and pKMDDV_Plasmid_2a (Addgene plasmid ID: 249056) as backbones. The msfGFP sequence was a gift from Michael Laub. The NahR and TetR sequences were taken from pAJM.771 (Addgene plasmid ID: 108534) and pAJM.011 (Addgene plasmid ID: 108529), respectively. DNA oligos and genes were ordered using the Oligo Rapid Service from Azenta Life Sciences. DNA fragments to be assembled were amplified by PCR using Phusion High-Fidelity PCR Master Mix with GC Buffer (NEB, M0532S). The PCR product underwent gel electrophoresis for 30 minutes at 110 V, as an additional quality control measure. Zymoclean Gel DNA Recovery Kit (Zymo Research, D4002) was used for gel extraction. Later, the plasmid is assembled with Gibson assembly protocol using Gibson Assembly Master Mix (NEB, E2611S). Assembled DNA was transformed into NEB5*α* (NEB, C2987H) for plasmid amplification. Transformed cells are then plated on Agar plates with 1.2% of Difco Agar and 2% of LB broth Lennox and appropriate antibiotics and grown overnight in an incubator at 37°C. Overnight cultures of singular colonies picked from the agar plates are grown in culture tubes with LB broth Lennox at 37°C. Plasmid DNA is then extracted from the overnight cultures using the plasmid miniprep-classic kit (Zymo Research, D4015). DNA sequencing is outsourced and performed by Plasmidsaurus: Standard Low Concentration Plasmid sequencing. After plasmid extraction from NEB5*α* and sequence confirmation, plasmids were transformed into MG1655 competent cells prepared by using CCMB80 buffer according to the manufacturer’s instructions (TekNova, C3132). The lists of plasmids and genetic parts are listed in Tables S7.

### Sample preparation for temporal measurements

Measurement was taken by briefly vortexing the bacterial culture, then withdrawing 50 uL of culture (500 uL for the 3 hour timepoint) into a separate 1.5 mL tube. From this tube, 5 uL were used for flow cytometry and absolute cell counting, 10 uL were used for plasmid copy number quantification, and the remaining volume was pelleted and stored for RNA extraction.

### Flow cytometry and absolute cell counting

Bacterial samples were prepared for flow cytometry by diluting cells 100x in a solution of 0.145 M NaCl, 30 uM propidium iodide (Invitrogen, P1304MP), and 10^6^ microspheres/mL (Invitrogen, B7277). Flow cytometry was then performed using a BD FACS Symphony A1.

The number of live (*n*_*live*_) and dead (*n*_*dead*_) cells per mL culture were counted by dividing the number of PI+ (*E*_*PI*−_) and PI-(*E*_*PI*+_) cell events, respectively, against the number of microsphere events (*E*_*MS*_), then multiplying by a dilution factor, *d* = 10^8^, using the below formulae.

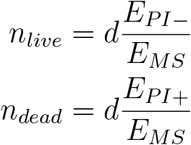

### Plasmid copy number quantification

Plasmid copy number was computed using the ΔΔ*C*_*q*_ method as in [49]. PowerUp™ SYBR™ Green Master Mix was used to perform qPCR (Applied Biosystems, A25918).

### RNA Extraction

Bacterial culture was pelleted (16000 rcf for 3 min.) and supernatant removed before storing at −80°C for no longer than 2 weeks, until performing RNA extraction. Total RNA was extracted using the Monarch Spin RNA Isolation Kit (NEB, T2110L) according to the manufacturer’s instructions. Specifically, the NEBExpress® T4 Lysozyme (NEB, P8115L) was used, also according to manufacturer’s instructions, prior to using Monarch StabiLyse™ DNA/RNA Buffer to ensure efficient cell lysis of stationary phase cells. The optional DNAseI treatment step was performed using DNase I-XT (NEB, M0570L).

### Estimation of dead cell half life

1 mL of an overnight culture of bacteria was pelleted (10000 rcf for 3 min.) and resuspended in 70% ethanol. After incubating at room temperature for 30 minutes, the cells were pelleted and washed with 0.85% NaCl before pelleting again and resuspending in M9 minimal media. Samples were transferred to a shaking incubator (30°C and 250 rpm). Flow cytometry measurements were taken from 0 – 168 hours in 24 hour increments. Half life *t*_1*/*2_ of PI+ cells was estimated using a best fit to the following equation:

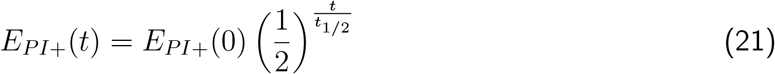

### Absolute RNA quantification

To obtain a mapping between the absolute number of cDNA and C_q_, a standard curve was generated using a plasmid-borne version of the msfGFP fragment being targeted for qPCR. Absolute counts of the plasmid were computed according to [50], then diluted to concentrations ranging across 5 orders of magnitude (10^4^ − 10^8^ copies/uL). qPCR using Luna® Universal qPCR Master Mix (NEB, M3003S) was performed on each concentration in technical triplicates to produce a standard curve.

msfGFP RNA was quantified in purified RNA samples through reverse transcriptase-qPCR (RT-qPCR) using the Luna® Universal One-Step RT-qPCR Kit (NEB, E3005L) on a Roche LightCycler® 96. The obtained C_q_ was first mapped back to the absolute copy number through the previously-generated standard curve, then this value was used to compute the number of RNA counts per RNA extraction. Because absolute cell count was also measured (see above), the RNA counts per cell is computed as the total msfGFP RNA count per RNA extraction divided by the total cell count per RNA extraction.

### Exponential-to-stationary phase model fitting and selection routine for characterization experiments

Model parameters were estimated using SciPy’s ‘curve_fit’ function with the Levenberg–Marquardt algorithm.^51^ Sample collection time and the mean of each species’ triplicate measurements were the independent and dependent variables, respectively. Weights for fitting each data point through the “sigma” argument were computed by accounting for both temporal sampling density and measurement variability using the formula

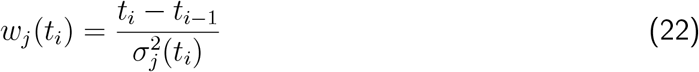

where *t*_*i*_ is the time of the *i*-th sampling time point out of *N* sampling time points (*i* ∈ *{*1…*N}*) and 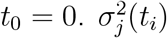 is the variance in measurement of species *j* measured at time *t*_*i*_.

Models were fit sequentially by first estimating the parameters associated with cell counts, then fixing cell count-associated parameters and fitting plasmid copy number. Then, parameters associated with both cell counts and PCN were fixed before fitting RNA copy number for each promoter:RBS pair. Finally, cell count, PCN, and RNA count parameters were fixed before fitting protein fluorescence values. However, protein fluorescence was fit by simultaneously optimizing one set of protein-associated parameters for each promoter:RBS pair sharing the same RBS.

Model selection was performed using residual sum of squares to compute the Akaike Information Criterion given by the following equations derived in [27]:

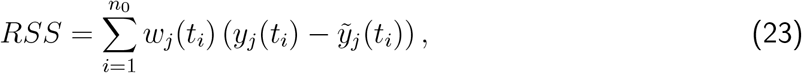

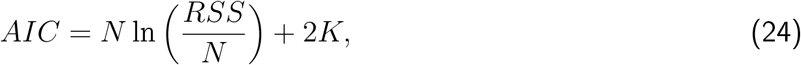

where *K* is the number of estimated parameters, *y*_*j*_(*t*_*i*_) is the average of the measured data at time 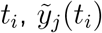 is the simulated value at time *t*_*i*_, and *n*_0_ is the number of timepoints.

## Supplementary Materials

**Table S1:** Predicted strengths of each characterized promoter and RBS using the De Novo DNA’s RBS calculator Version 2020 and Promoter calculator Version 2022.^1^.

| Type | Component | Strength |
| --- | --- | --- |
| Promoter | J23102 | 9749 |
| Promoter | J23119 | 35692 |
| RBS | B0034a | 2251 |
| RBS | J61119 | 38 |
| RBS | J61136 | 191 |

**Table S2:** Predicted strengths of the promoters and RBSs used to drive regulator expression using the De Novo DNA’s RBS calculator Version 2020 and Promoter calculator Version 2022.^1^.

| Type | Component | Strength |
| --- | --- | --- |
| Promoter | J23114 | 6065 |
| Promoter | J23102 | 2877 |
| RBS | $R_{rbs}$ (NahR) | 1375 |
| RBS | $R_{rbs}$ (TetR) | 10248 |

**Table S6:**
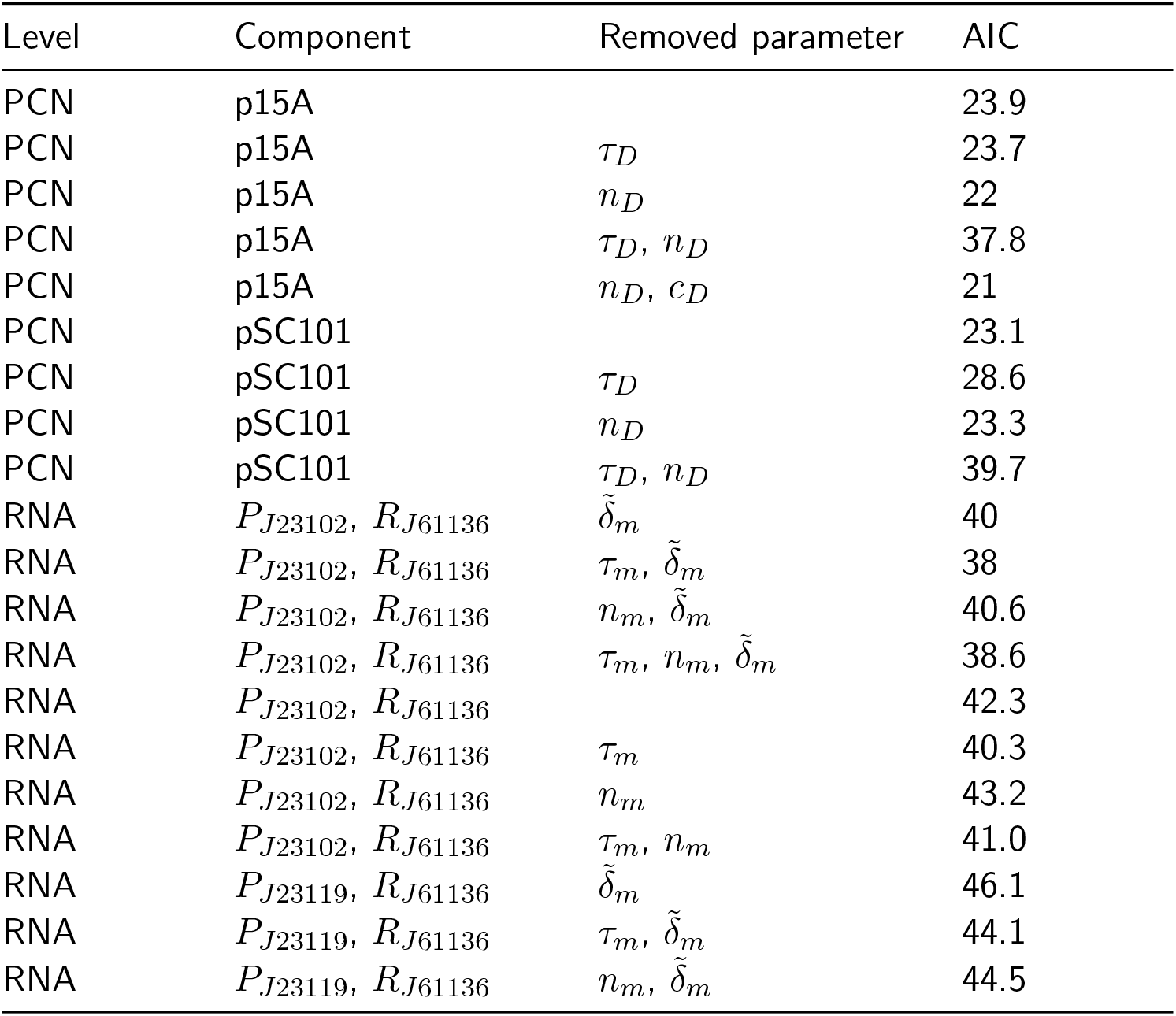
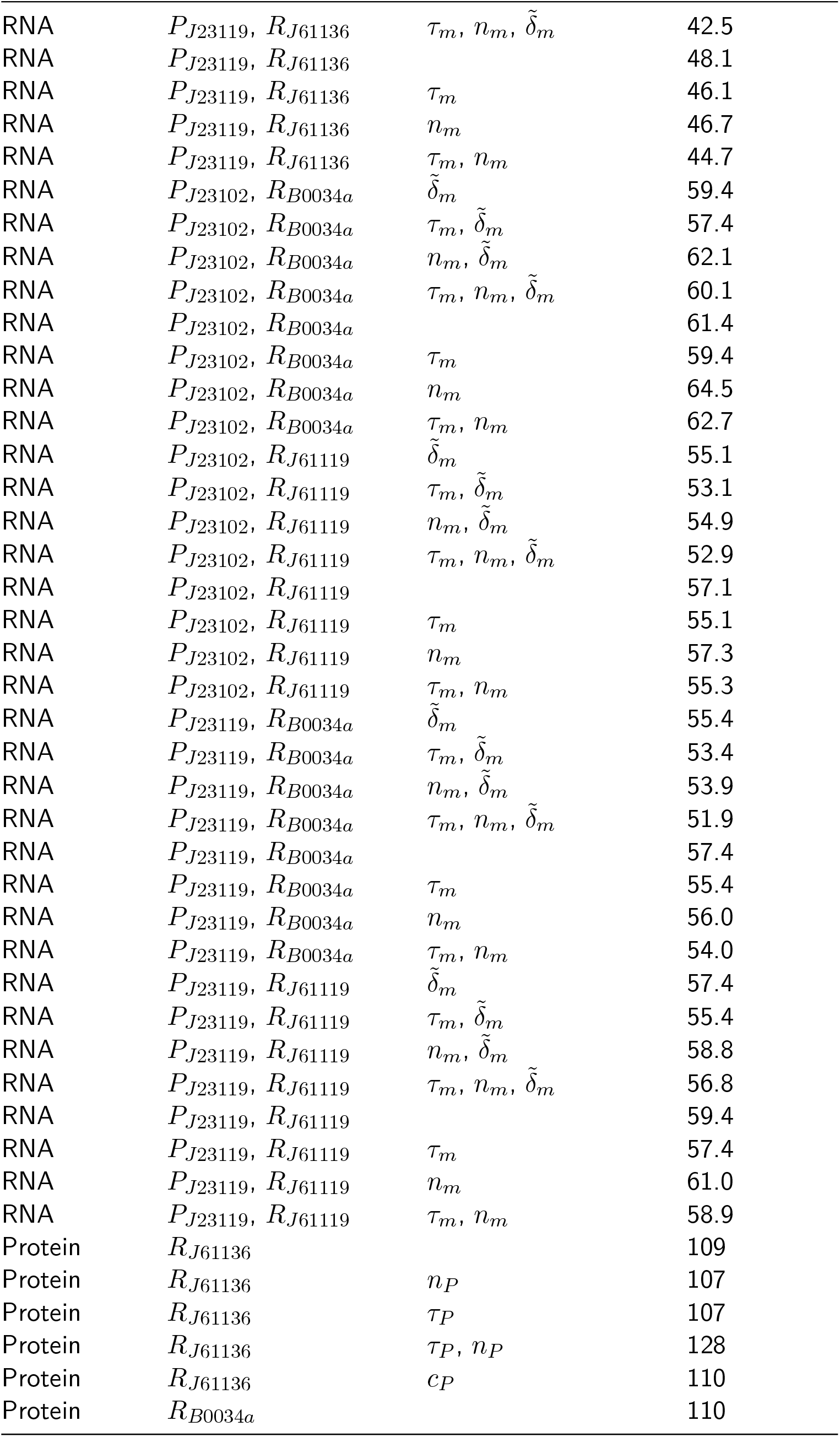
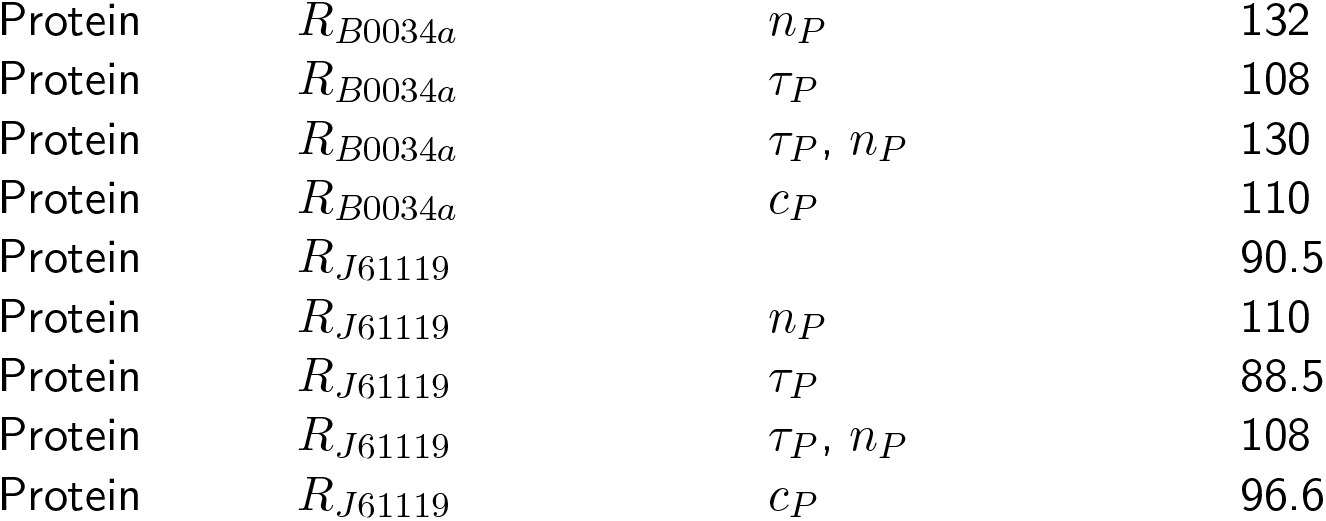
AIC values for model variants.

| Level | Component | Removed parameter | AIC |
| --- | --- | --- | --- |
| PCN | p15A |  | 23.9 |
| PCN | p15A | $\tau_D$ | 23.7 |
| PCN | p15A | $n_D$ | 22 |
| PCN | p15A | $\tau_D, n_D$ | 37.8 |
| PCN | p15A | $n_D, c_D$ | 21 |
| PCN | pSC101 |  | 23.1 |
| PCN | pSC101 | $\tau_D$ | 28.6 |
| PCN | pSC101 | $n_D$ | 23.3 |
| PCN | pSC101 | $\tau_D, n_D$ | 39.7 |
| RNA | $P_{J23102}, R_{J61136}$ | $\tilde{\delta}_m$ | 40 |
| RNA | $P_{J23102}, R_{J61136}$ | $\tau_m, \tilde{\delta}_m$ | 38 |
| RNA | $P_{J23102}, R_{J61136}$ | $n_m, \tilde{\delta}_m$ | 40.6 |
| RNA | $P_{J23102}, R_{J61136}$ | $\tau_m, n_m, \tilde{\delta}_m$ | 38.6 |
| RNA | $P_{J23102}, R_{J61136}$ | | 42.3 |
| RNA | $P_{J23102}, R_{J61136}$ | $\tau_m$ | 40.3 |
| RNA | $P_{J23102}, R_{J61136}$ | $n_m$ | 43.2 |
| RNA | $P_{J23102}, R_{J61136}$ | $\tau_m, n_m$ | 41.0 |
| RNA | $P_{J23119}, R_{J61136}$ | $\tilde{\delta}_m$ | 46.1 |
| RNA | $P_{J23119}, R_{J61136}$ | $\tau_m, \tilde{\delta}_m$ | 44.1 |
| RNA | $P_{J23119}, R_{J61136}$ | $n_m, \tilde{\delta}_m$ | 44.5 |
| RNA | $P_{J23119}, R_{J61136}$ | $\tau_m, n_m, \tilde{\delta}_m$ | 42.5 |
| RNA | $P_{J23119}, R_{J61136}$ | | 48.1 |
| RNA | $P_{J23119}, R_{J61136}$ | $\tau_m$ | 46.1 |
| RNA | $P_{J23119}, R_{J61136}$ | $n_m$ | 46.7 |
| RNA | $P_{J23119}, R_{J61136}$ | $\tau_m, n_m$ | 44.7 |
| RNA | $P_{J23102}, R_{B0034a}$ | $\tilde{\delta}_m$ | 59.4 |
| RNA | $P_{J23102}, R_{B0034a}$ | $\tau_m, \tilde{\delta}_m$ | 57.4 |
| RNA | $P_{J23102}, R_{B0034a}$ | $n_m, \tilde{\delta}_m$ | 62.1 |
| RNA | $P_{J23102}, R_{B0034a}$ | $\tau_m, n_m, \tilde{\delta}_m$ | 60.1 |
| RNA | $P_{J23102}, R_{B0034a}$ | | 61.4 |
| RNA | $P_{J23102}, R_{B0034a}$ | $\tau_m$ | 59.4 |
| RNA | $P_{J23102}, R_{B0034a}$ | $n_m$ | 64.5 |
| RNA | $P_{J23102}, R_{B0034a}$ | $\tau_m, n_m$ | 62.7 |
| RNA | $P_{J23102}, R_{J61119}$ | $\tilde{\delta}_m$ | 55.1 |
| RNA | $P_{J23102}, R_{J61119}$ | $\tau_m, \tilde{\delta}_m$ | 53.1 |
| RNA | $P_{J23102}, R_{J61119}$ | $n_m, \tilde{\delta}_m$ | 54.9 |
| RNA | $P_{J23102}, R_{J61119}$ | $\tau_m, n_m, \tilde{\delta}_m$ | 52.9 |
| RNA | $P_{J23102}, R_{J61119}$ | | 57.1 |
| RNA | $P_{J23102}, R_{J61119}$ | $\tau_m$ | 55.1 |
| RNA | $P_{J23102}, R_{J61119}$ | $n_m$ | 57.3 |
| RNA | $P_{J23102}, R_{J61119}$ | $\tau_m, n_m$ | 55.3 |
| RNA | $P_{J23119}, R_{B0034a}$ | $\tilde{\delta}_m$ | 55.4 |
| RNA | $P_{J23119}, R_{B0034a}$ | $\tau_m, \tilde{\delta}_m$ | 53.4 |
| RNA | $P_{J23119}, R_{B0034a}$ | $n_m, \tilde{\delta}_m$ | 53.9 |
| RNA | $P_{J23119}, R_{B0034a}$ | $\tau_m, n_m, \tilde{\delta}_m$ | 51.9 |
| RNA | $P_{J23119}, R_{B0034a}$ | | 57.4 |
| RNA | $P_{J23119}, R_{B0034a}$ | $\tau_m$ | 55.4 |
| RNA | $P_{J23119}, R_{B0034a}$ | $n_m$ | 56.0 |
| RNA | $P_{J23119}, R_{B0034a}$ | $\tau_m, n_m$ | 54.0 |
| RNA | $P_{J23119}, R_{J61119}$ | $\tilde{\delta}_m$ | 57.4 |
| RNA | $P_{J23119}, R_{J61119}$ | $\tau_m, \tilde{\delta}_m$ | 55.4 |
| RNA | $P_{J23119}, R_{J61119}$ | $n_m, \tilde{\delta}_m$ | 58.8 |
| RNA | $P_{J23119}, R_{J61119}$ | $\tau_m, n_m, \tilde{\delta}_m$ | 56.8 |
| RNA | $P_{J23119}, R_{J61119}$ | | 59.4 |
| RNA | $P_{J23119}, R_{J61119}$ | $\tau_m$ | 57.4 |
| RNA | $P_{J23119}, R_{J61119}$ | $n_m$ | 61.0 |
| RNA | $P_{J23119}, R_{J61119}$ | $\tau_m, n_m$ | 58.9 |
| Protein | $R_{J61136}$ | | 109 |
| Protein | $R_{J61136}$ | $n_P$ | 107 |
| Protein | $R_{J61136}$ | $\tau_P$ | 107 |
| Protein | $R_{J61136}$ | $\tau_P, n_P$ | 128 |
| Protein | $R_{J61136}$ | $c_P$ | 110 |
| Protein | $R_{B0034a}$ | | 110 |
| Protein | $R_{B0034a}$ | $n_P$ | 132 |
| Protein | $R_{B0034a}$ | $\tau_P$ | 108 |
| Protein | $R_{B0034a}$ | $\tau_P, n_P$ | 130 |
| Protein | $R_{B0034a}$ | $c_P$ | 110 |
| Protein | $R_{J61119}$ | | 90.5 |
| Protein | $R_{J61119}$ | $n_P$ | 110 |
| Protein | $R_{J61119}$ | $\tau_P$ | 88.5 |
| Protein | $R_{J61119}$ | $\tau_P, n_P$ | 108 |
| Protein | $R_{J61119}$ | $c_P$ | 96.6 |

**Table S7:**
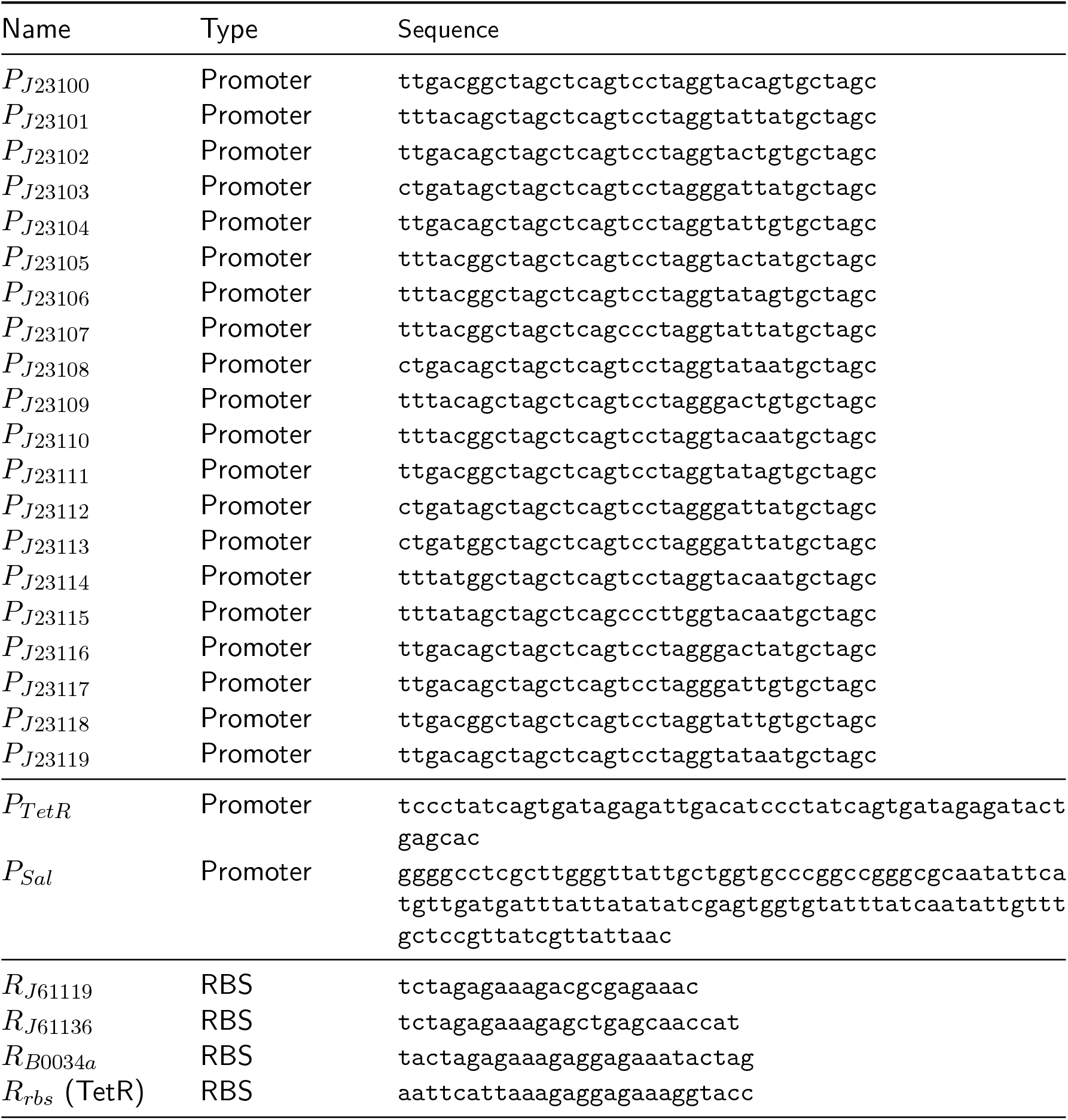
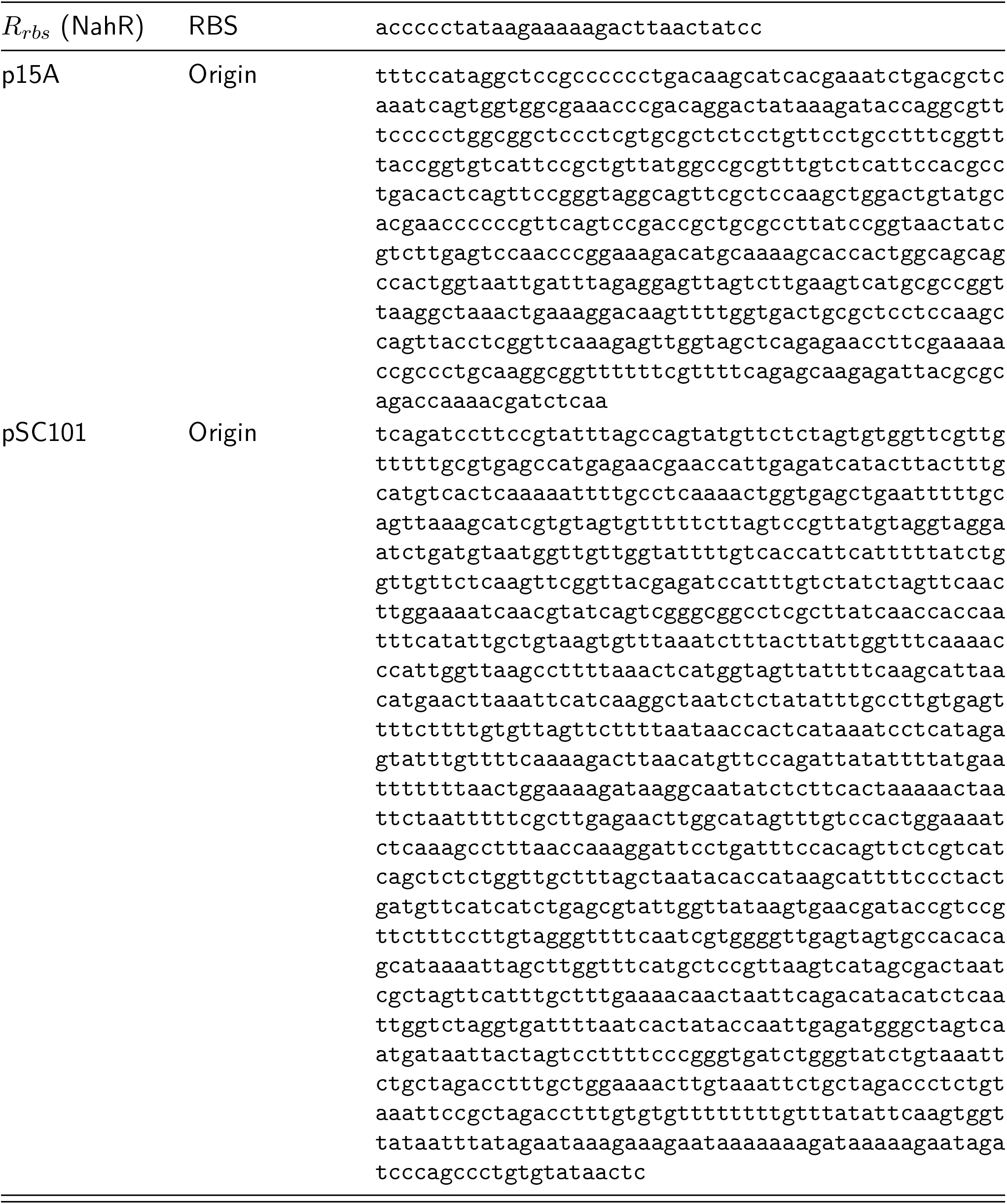
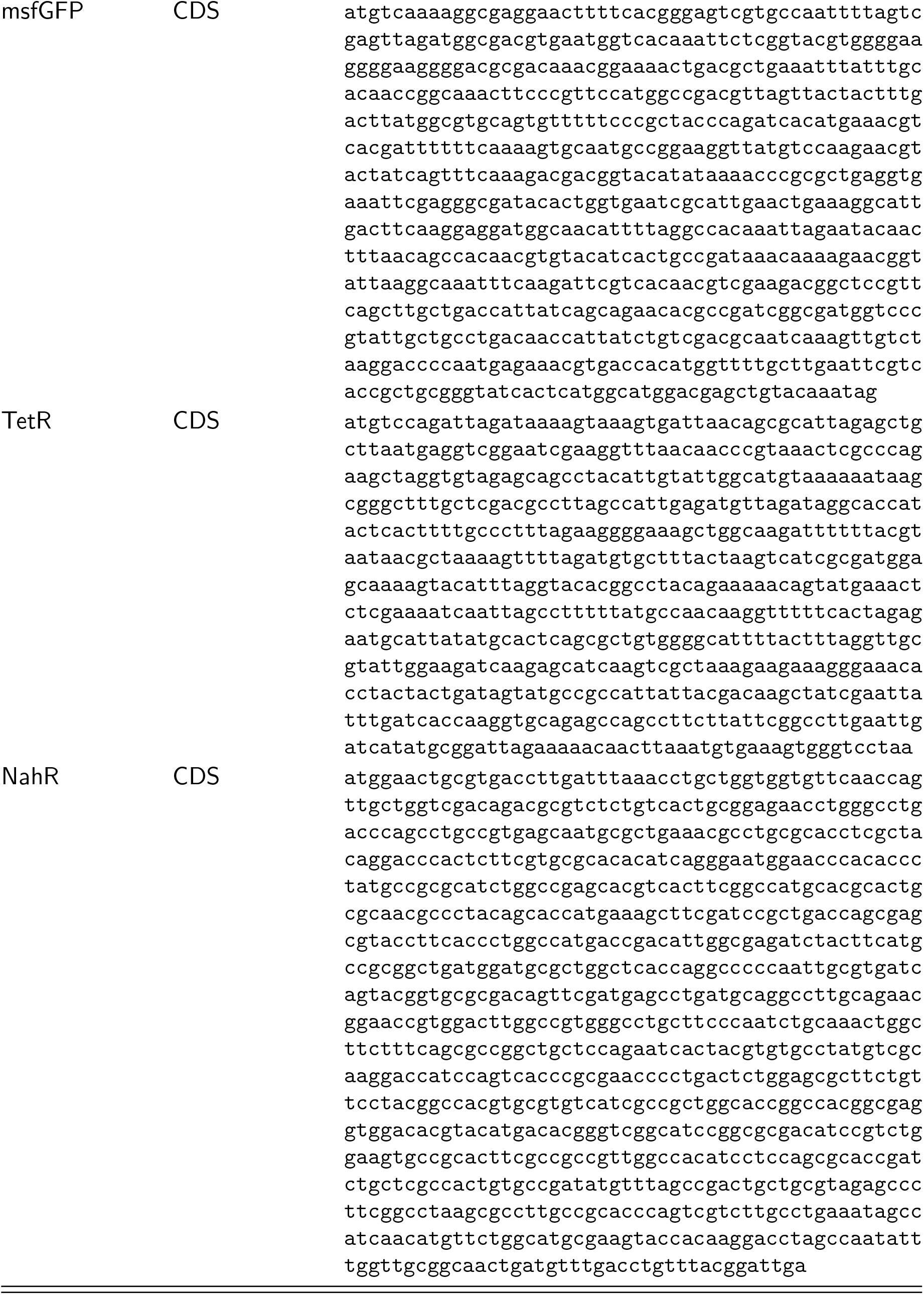
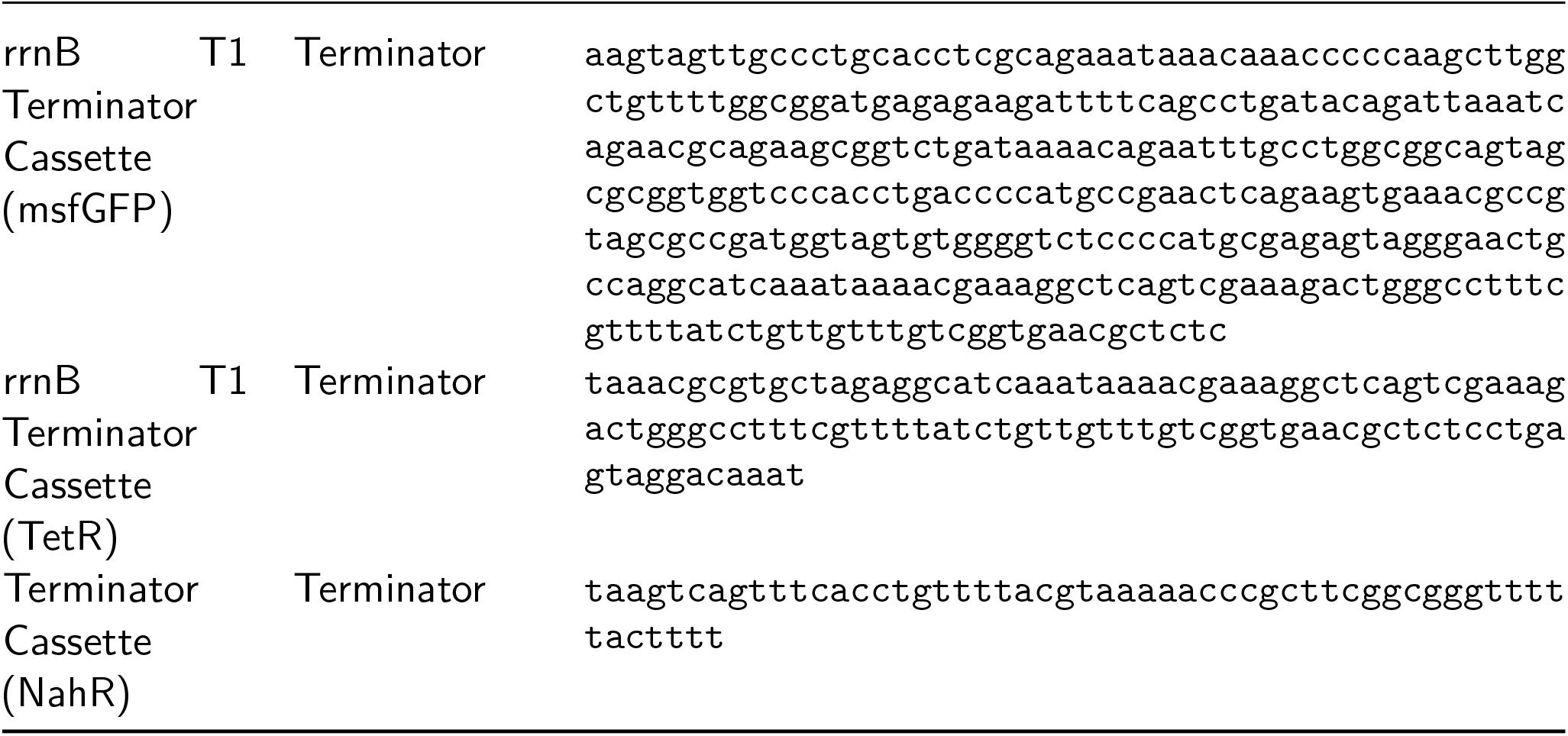
DNA sequences for genetic parts.

**Fig. S1.**
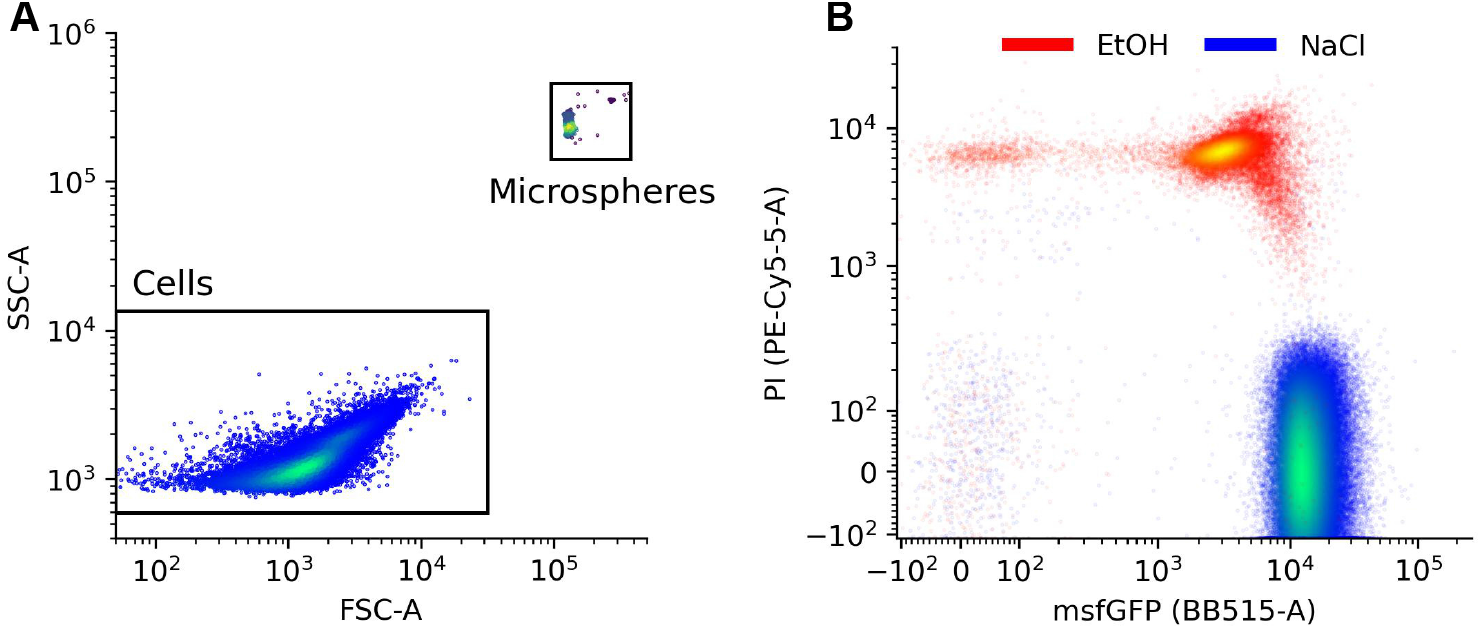
Absolute live/dead cell count measurements. (A) Flow cytometry measurements of a sample of cells and microspheres. Number of cells are normalized by the number of microspheres to compute cells/mL (Methods). (B) Flow cytometry measurements of cells harboring a constitutively-expressed msfGFP construct after being incubated in 70% Ethanol (EtOH) or a 0.85% saline (Methods). 70% ethanol is known to cause cell death by compromising bacterial cell membranes.^2^

**Fig. S2.**
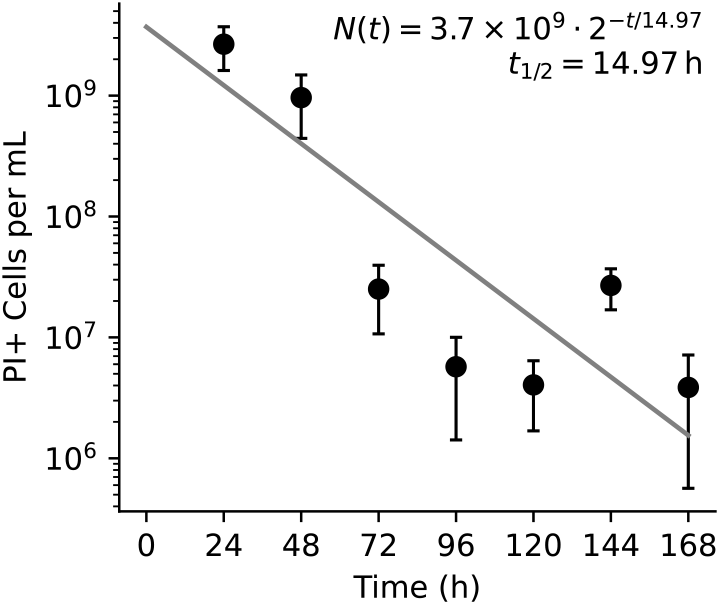
Measurement of PI+ cells over a week in culture after incubation in 70% EtOH to used to estimate PI+ cells’ half-life (see Methods). Solid line: OLS linear regression best fit of the decaying exponential indicated in the plot. Data points in each panel represent the mean ± one standard deviation calculated from three biological replicates.

**Fig. S3.**
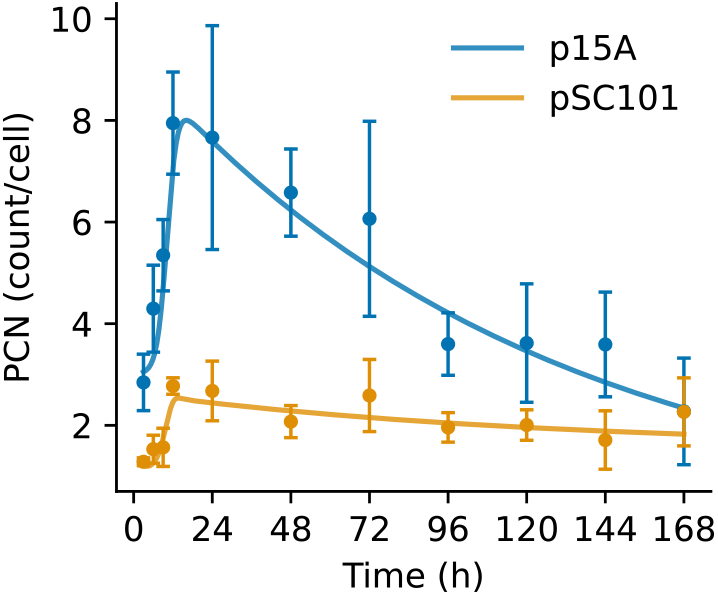
Plasmid copy number over time for both p15A and pSC101 origins of replication. Solid line: OLS best fit of Eq. 5 based on the data shown (see Methods; parameters in Table S4)

**Fig. S4.**
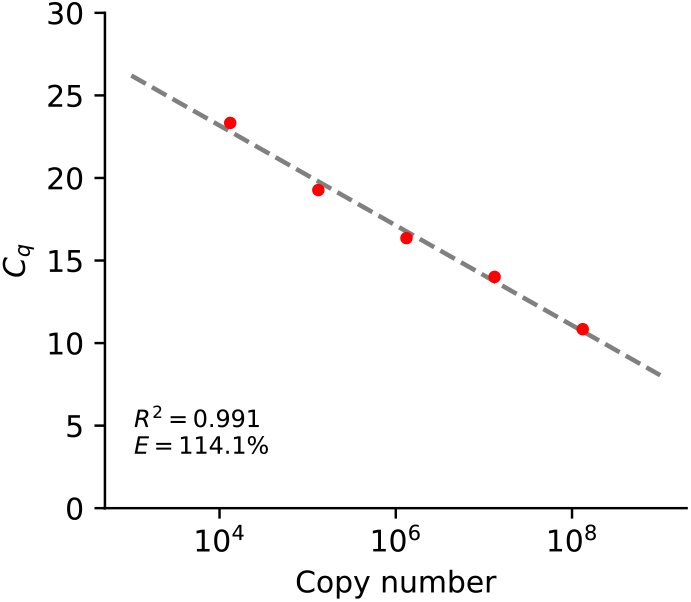
Standard curve generated for absolute quantification of msfGFP RNA.

**Fig. S5.**
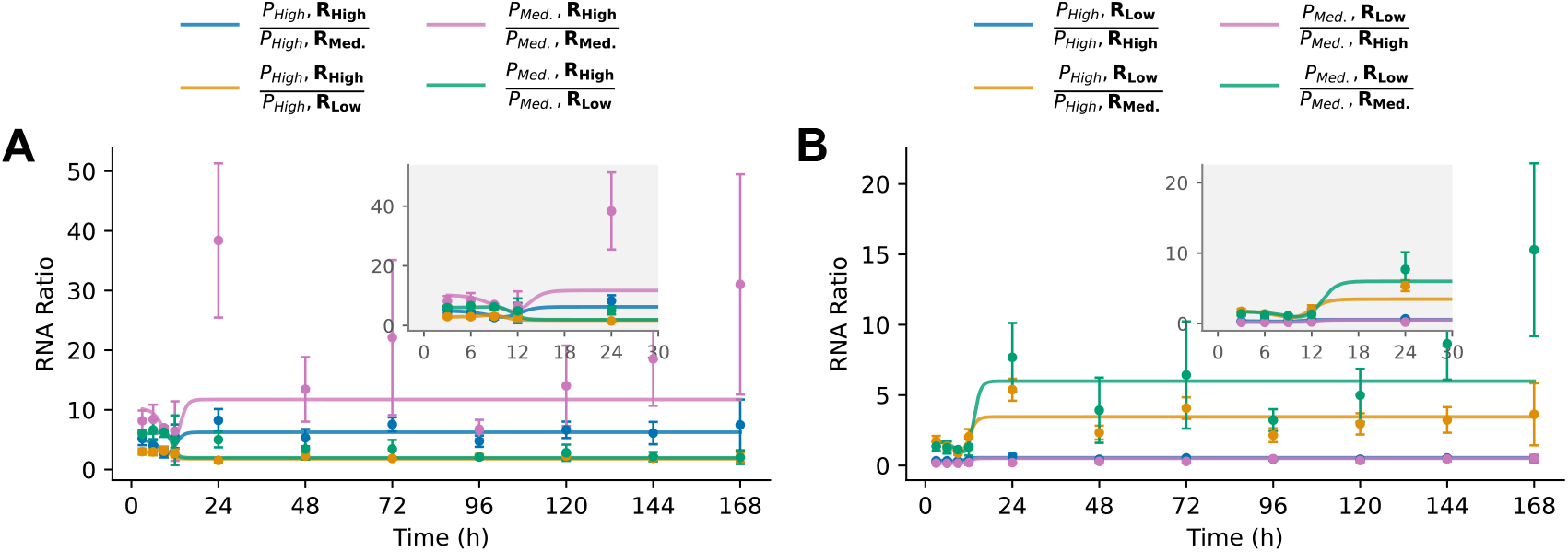
(A) Ratio of RNA counts between constructs harboring the high-strength RBS *R*_*High*_ (numerator) and each other construct as the denominator: *P*_*High*_ with medium-strength RBS *R*_*Med*._ (blue), *P*_*High*_ with low-strength RBS *R*_*Low*_ (orange), *P*_*Med*._ with medium-strength RBS *R*_*Med*._ (pink), *P*_*Med*._ with low-strength RBS *R*_*Low*_ (green). (B) Ratio of RNA counts between constructs harboring the low-strength RBS *R*_*Low*_ (numerator) and each other construct as the denominator: *P*_*High*_ with high-strength RBS *R*_*High*_ (blue), *P*_*High*_ with medium-strength RBS *R*_*Med*._ (orange), *P*_*Med*._ with high-strength RBS *R*_*High*_ (pink), *P*_*Med*._ with medium-strength RBS *R*_*Med*._ (green). Comparison groups are bolded in the legend for clarity. Solid lines: transcript ratios based on simulated OLS best fits seen in panel 3B.

**Fig. S6.**
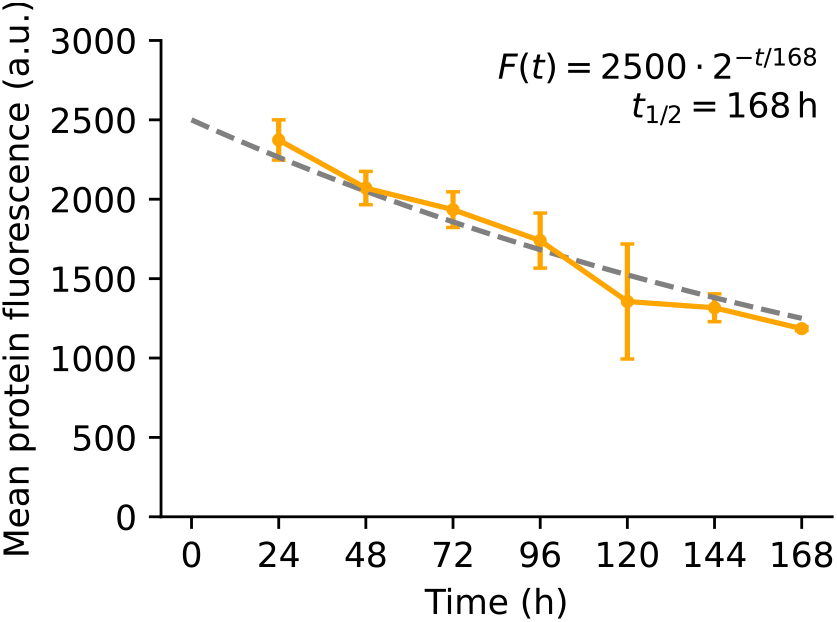
Measurement of msfGFP decay over time after 24-hours of applying aTc to the TetR system seen in Figure 5D. Thereafter, cells were washed with saline and monitored for the the remainder of the week. Dashed line represents the model of protein half-life. Half-life coarsely estimated by hand.

**Fig. S7.**
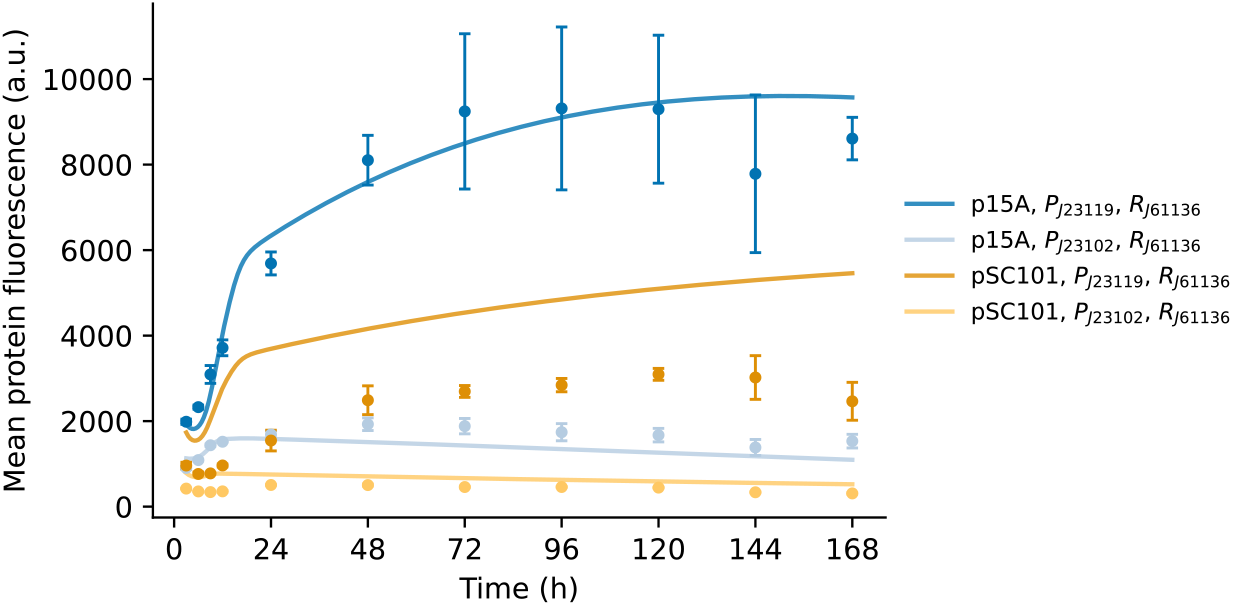
Protein fluorescence intensity over time for both p15A and pSC101 origins of replication. Solid blue lines: OLS best fit of combinations with p15A origin of replication. Solid orange lines: predicted protein fluorescence intensities based on the OLS best fit of the pSC101 origin of replication (see Methods; parameters in Table S4), allowing all other RNA count and protein fluorescence parameters fixed as those estimated from combinations using the p15A origin of replication.

**Table S3:** Parameter estimates associated with the inducible promoter systems in Eq. 9–19.

| Parameter | Value | Unit |
| --- | --- | --- |
| $\alpha_N$ | 0.719 | $\text{h}^{-1}$ |
| $K_N$ | $3.81 \times 10^9$ | $\text{Cells } \text{mL}^{-1}$ |
| $\tau_D$ | 3.60 | $\text{h}$ |
| $\alpha_D$ | 2.20 | $\text{PCN}^{-1} \text{h}^{-1}$ |
| $\delta_D$ | 0.00815 | $\text{h}^{-1}$ |
| $\alpha_N$ | 0.719 | $\text{h}^{-1}$ |
| $\delta_m$ | 8.32 | $\text{h}^{-1}$ |
| $n_m(P_{Sal})$ | 1.51 | - |
| $n_m(P_{TetR})$ | 1.2 | - |
| $c_m(P_{Sal})$ | $1.48 \times 10^{-3}$ | - |
| $c_m(P_{TetR})$ | $4.65 \times 10^{-4}$ | - |
| $\alpha_m(P_{Sal})$ | 1590 | $\text{h}^{-1}$ |
| $\alpha_m(P_{TetR})$ | 1910 | $\text{h}^{-1}$ |
| $n_{m,A}$ | 1.45 | - |
| $n_{m,R}$ | 1.45 | - |
| $c_{m,A}$ | $4.20 \times 10^{-3}$ | - |
| $c_{m,R}$ | $4.20 \times 10^{-3}$ | - |
| $\alpha_{m,A}$ | 122 | $\text{h}^{-1}$ |
| $\alpha_{m,R}$ | 122 | $\text{h}^{-1}$ |
| $\tau_P$ | 3.60 | $\text{h}$ |
| $\alpha_P$ | 2.20 | $\text{PCN}^{-1} \text{h}^{-1}$ |
| $K_m$ | 30.6 | **a.u.** |
| $c_P$ | 0.719 | - |
| $\tau_A$ | 5.80 | $\text{h}$ |
| $\alpha_A$ | $1.11 \times 10^5$ | $\text{PCN}^{-1} \text{h}^{-1}$ |
| $K_{m,A}$ | $1.56 \times 10^3$ | **a.u.** |
| $c_A$ | 0.114 | - |
| $\tau_R$ | 5.80 | $\text{h}$ |
| $\alpha_R$ | $1.11 \times 10^5$ | $\text{PCN}^{-1} \text{h}^{-1}$ |
| $K_{m,R}$ | $1.56 \times 10^3$ | **a.u.** |
| $c_R$ | 0.114 | - |
| $\delta_P$ | $4.2 \times 10^{-3}$ | $\text{h}^{-1}$ |

**Table S4:** OLS parameter estimates associated with the pSC101 replication origins from the full PCN model in Eq. 2.

| Parameter | Value | Unit |
| --- | --- | --- |
| $\tau_D$ | 3.60 | $\text{h}$ |
| $\alpha_D$ | 0.816 | $\text{PCN}^{-1} \text{h}^{-1}$ |
| $n_D$ | 3.154 | - |
| $c_D$ | 0.0155 | - |

**Table S5:** OLS parameter estimates for each promoter in the Anderson Library, excluding *P*_*J*23102_ and *P*_*J*23119_ using the reduced RNA model in Eq. 7 with *n*_*m*_ = 1.43 fixed. *α*_*m*_ in *RNA/*(*PCN* · *h*).

| Promoter | $c_m$ | $\alpha_m$ | $n_m$ |
| --- | --- | --- | --- |
| $P_{J23100}$ | 0.0186 | 84.6 | 1.74 |
| $P_{J23101}$ | 0.00981 | 121 | 1.43 |
| $P_{J23103}$ | 0.0137 | 3.07 | 2.80 |
| $P_{J23104}$ | 0.00199 | 785 | 1.50 |
| $P_{J23105}$ | 0.0353 | 9.58 | 1.99 |
| $P_{J23106}$ | 0.0167 | 18.4 | 1.42 |
| $P_{J23107}$ | 0.0133 | 9.31 | 1.32 |
| $P_{J23108}$ | 0.0185 | 24.5 | 2.18 |
| $P_{J23109}$ | 0.0215 | 3.55 | 2.30 |
| $P_{J23110}$ | 0.00592 | 33.7 | 1.46 |
| $P_{J23111}$ | 0.00115 | 4900 | 3.51 |
| $P_{J23112}$ | 0.0137 | 3.15 | 2.70 |
| $P_{J23113}$ | 0.0105 | 3.49 | 2.75 |
| $P_{J23114}$ | 0.0139 | 3.84 | 2.44 |
| $P_{J23115}$ | 0.0143 | 5.58 | 1.97 |
| $P_{J23116}$ | 0.0192 | 4.32 | 1.53 |
| $P_{J23117}$ | 0.0202 | 3.22 | 2.09 |
| $P_{J23118}$ | 0.00622 | 107 | 1.07 |

### Supplementary Note: Derivation of regulator complexes

The activator complex *C*_*A*_ in Eq. 13 and free repressor *R* in Eq. 19 are derived from equations describing the regulators when free and bound to their respective small molecule inducer. Beginning with the NahR (*A*) activation system, we model the dynamics of the free and unbound forms of *A* as follows:

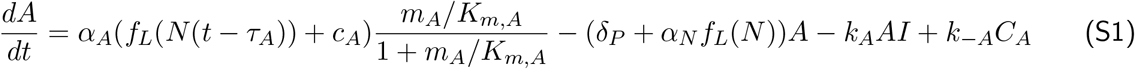

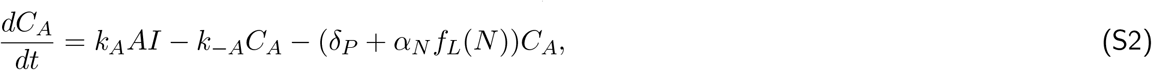

where *A* is the free activator protein *A, I* is the small molecule inducer, and *C*_*A*_ is the complex of *A* bound with *I*. Using the quasi-steady state approximation to set 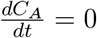 and applying the conservation law *A*_*T*_ = *A* + *C*_*A*_, we have:

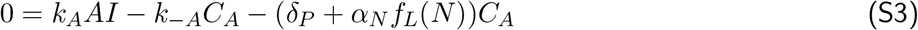

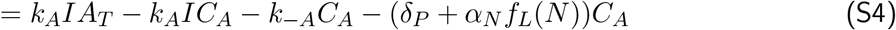

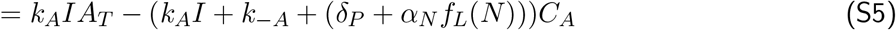

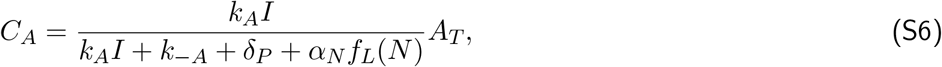

since *δ*_*P*_ + *α*_*N*_ *f*_*L*_(*N*) ≪ *k*_− *A*_, given that reversible binding dynamics are typically much faster than molecule decay.^3^ Therefore:

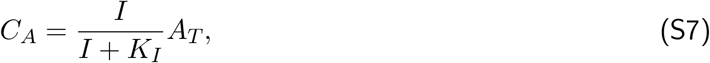

with 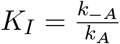. By writing the differential equation for *A*_*T*_, we further have:

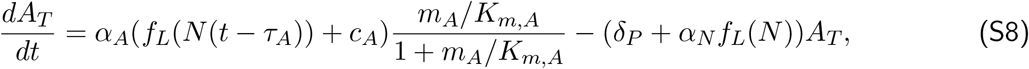

which is the same expression as Eq. 13, in which *A* is to be intended as *A*_*T*_.

We use a similar approach to derive the free amount of repressor protein *R*_*free*_ in Eq. 19 and total repressor *R*_*T*_ (referred to as *R* in the main text) in Eq. 16:

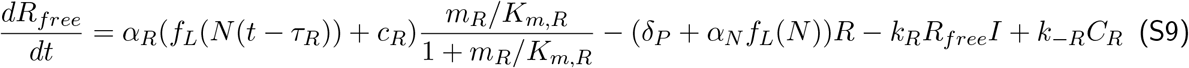

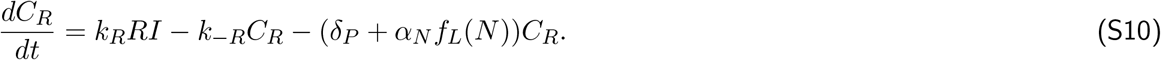

We set 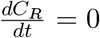, apply the conservation law *R*_*T*_ = *R* + *R*_*free*_, and account for the fact that the reversible binding dynamics of are much faster than molecule decay:

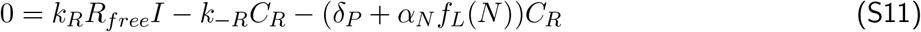

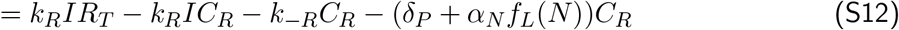

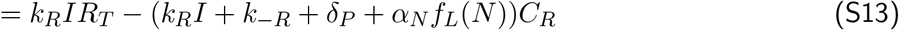

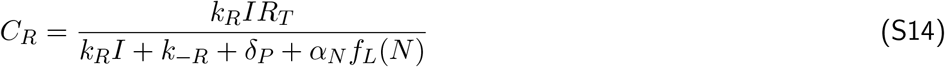

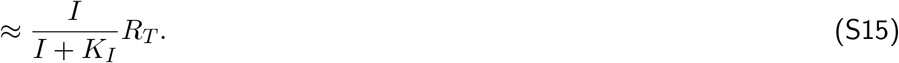

Then, because *R*_*free*_ = *R*_*T*_ − *C*_*R*_:

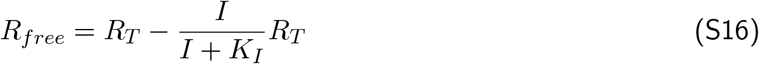

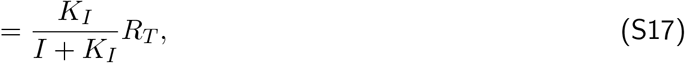

which is the same expression as Eq. 19, in which *R* is intended as *R*_*T*_.

Derivations for the activated promoter *D*_*A*_ (*C*_*I*_ bound to *D*) in Eq. 14, and the free promoter *D*_*free*_ in Eq. 20 follow a similar process. Beginning with *D*_*A*_, we have the following differential equations:

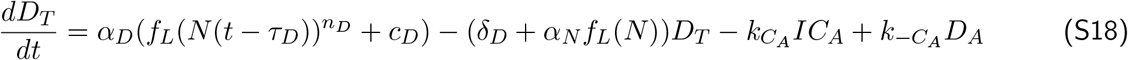

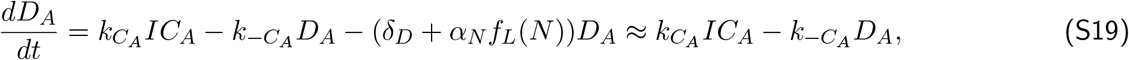

where *D*_*T*_ is the total plasmid copy number.

Applying the conservation law *D*_*T*_ = *D*_*A*_ + *D*_*free*_ and using the quasi-steady state approximation to set 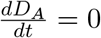:

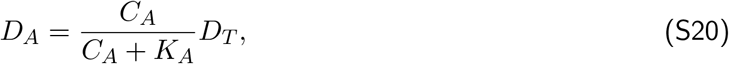

which is the same differential equation as Eq. 14, in which D is intended as *D*_*T*_.

Similarly, to derive *D*_*free*_ from Eq. 20, we begin with the following differential equations:

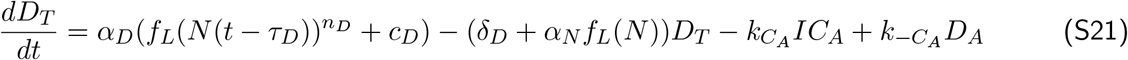

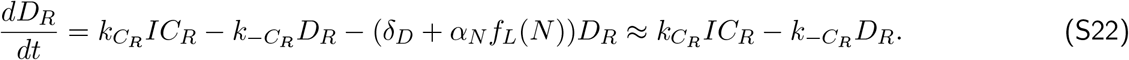

Applying the conservation law *D*_*T*_ = *D*_*R*_ + *D*_*free*_ and using the quasi-steady state approximation to set 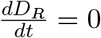 yields:

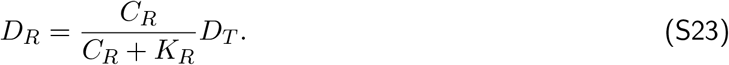

Then, solving for *D*_*free*_ given that *D*_*free*_ = *D*_*T*_ − *D*_*R*_ yields:

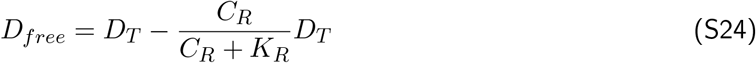

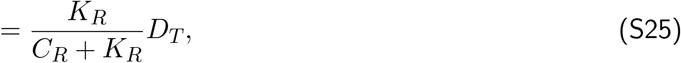

which is the same differential equation as Eq. 20, in which D is intended as *D*_*T*_.

